# The MIRO1-BAX Complex Dictates Life and Death at the Mitochondrial Gate

**DOI:** 10.64898/2026.06.29.733391

**Authors:** Alva G. Sainz, Chulhwan S. Kwak, Kwang Bog Cho, Sejal A. Sripadanna, Brandon H. Bergsneider, Zoe Zizzo, Aarooran S. Durairaj, Zehui Du, Ian Cooney, Anthony Venida, Nike Bharucha, Ioannis Karakikes, Wah Chiu, Michael Lim, Michael C. Bassik, Xinnan Wang

## Abstract

BAX macropores in the outer mitochondrial membrane (OMM) are canonical mediators of apoptosis, but whether the same pore structure can drive distinct cell death pathways remains unclear. Here, we identify the OMM protein MIRO1 as a context-specific modulator of BAX activity. Mechanistically, MIRO1 binds BAX via MIRO1’s N-terminal domain to promote macropore formation and the release of mitochondrial DNA (mtDNA) into the cytoplasm, triggering the STING-pIRF3 signaling axis. In glioma cells, this pathway sustains GPX4 expression via pIRF3-mediated transcriptional activation and confers ferroptosis resistance while bypassing inflammation. By contrast, in Parkinsonian neurons, the MIRO1-BAX complex promotes mitochondrial-stress-induced apoptosis. Using structure-guided drug discovery, we developed first-in-class small molecules that allosterically disrupt the MIRO1-BAX complex by engaging MIRO1’s distal GTPase pocket. These compounds sensitize glioma cells to ferroptosis and protect neurons from apoptosis. Our findings reveal a disease-specific mitochondrial switch for life-death decisions and illuminate the molecular logic by which cells exploit and interpret OMM permeabilization.

## Introduction

Mitochondria act as key signaling hubs (*1*) that integrate various death signals and promote cell death, in part, by releasing pro-death factors into the cytosol via distinct pores on the outer mitochondrial membrane (OMM). The composition of these pores varies with the type of cell death. For example, BAX macropores form during apoptotic or sub-apoptotic stress (*2*), whereas Gasdermin D (GSDMD) pores form during pyroptosis (*3*). BAX macropores can leak mitochondrial DNA (mtDNA) into the cytosol, activating the cGAS-STING pathway and downstream innate immune signaling (*2, 4–7*). While emerging evidence suggests crosstalk between distinct cell death programs (*8, 9*), it remains unclear whether a single pore structure, such as the BAX macropore, can mediate mechanistically divergent cell death pathways in a disease-specific manner. In addition to pro-apoptotic factors, OMM pores have been shown to mediate the release of other signaling molecules, including mtDNA, into the cytosol, triggering responses beyond canonical cell death programs, such as innate immune signaling (*2, 4–7*). Whether these additional signaling modalities triggered by OMM pores play a role in dictating cell death mechanisms in different disease contexts remains to be elucidated.

MIRO1 (*RHOT1*) is an OMM protein that regulates key mitochondrial behaviors, including mitochondrial transport (*10–12*) and mitophagy (*13, 14*). Anchored to the OMM via a C-terminal transmembrane (TM) domain, MIRO1 projects an N-terminal cytosolic region containing two EF-hand Ca²⁺-binding motifs and two GTPase domains. Its C-terminal tail extends into the mitochondrial intermembrane space (IMS). MIRO1’s diverse roles depend on dynamic protein–protein interactions: for mitochondrial trafficking, MIRO1 binds to the TRAK/kinesin (KHC) complex (*10–12*); during mitophagy signaling, it transiently associates with PINK1, Parkin, and LRRK2 before its proteasomal degradation (*13, 14*). MIRO1 can assemble into higher-order super-complexes (*15*), and its conformation and stoichiometry are modulated by post-translational modifications and cellular signals, including Ca²⁺, glucose, and hypoxia (*12, 13, 16*). These context-dependent changes underline MIRO1’s functional plasticity across cell types and physiological states.

Parkinson’s disease (PD) is a debilitating age-dependent neurodegenerative disease primarily affecting midbrain dopaminergic (DA) neurons. We have previously found that MIRO1 protein is upregulated in postmortem brains of sporadic PD patients and induced pluripotent stem cell (iPSC)-derived neurons of familial PD patients carrying *SNCA* mutations (*15, 17*). Stabilization of MIRO1 delays its removal from damaged mitochondria, postpones mitophagy, and exacerbates oxidative stress, increasing neuronal vulnerability to stressors in PD patient-derived models (*14, 15, 17–20*). To target MIRO1, we developed a series of compounds ("MIRO1 Reducers") using an artificial intelligence (AI)-guided virtual screen and functional tests in Drosophila and human cells, identifying candidates that bind to the C-terminal GTPase domain of human MIRO1 (*18, 19*). In PD patient fibroblasts and iPSC-derived neurons, these compounds at the therapeutic dose selectively promote proteasome-mediated degradation of MIRO1 upon mitochondrial depolarization, without affecting MIRO1’s overall GTPase activity, other mitochondrial proteins, or basal mitochondrial transport (*15, 18, 19*). MIRO1 Reducers rescue delayed mitophagy and protect against stress-induced DA neuron loss in PD models, highlighting MIRO1 as a promising therapeutic target (*15, 18, 19*).

Here, we uncover a previously unrecognized function of MIRO1: promoting BAX macropore formation at the OMM and dictating cell survival. MIRO1 forms a complex with BAX via MIRO1’s N-terminal domain, and disrupting this interaction blunts BAX activation and downstream signaling. Unexpectedly, this MIRO1-BAX complex has opposing effects across disease contexts. It facilitates mtDNA release and activates STING-pIRF3 to sustain GPX4 expression and confer resistance to ferroptosis in glioma cells. Conversely, it triggers apoptosis in PD-relevant neurons under oxidative stress. Through multi-scale drug screening and structural analyses, we identify new MIRO1 ligands with distinct physicochemical properties, providing a chemical toolkit for mechanistic studies across various disease models.

## Results

### *The MIRO1* gene is expressed in a higher proportion of cells in both PD and glioma brain tissue

Given MIRO1’s essential role in neuronal mitochondrial homeostasis, we examined its gene expression at the single-cell level in human brain tissue from patients with neurodegenerative or neoplastic diseases. We analyzed single-cell RNA-sequencing (scRNA-seq) data from (*21*), which includes 22,048 DA neurons from 8 neurotypical donors (healthy controls), 4 patients with Lewy Body Dementia (LBD), and 6 patients with PD (Fig. 1A). *MIRO1* was detected across all DA neuron subclusters and was expressed in a higher percentage of DA neurons from PD patients compared to both neurotypical and LBD controls (Fig. 1B). To evaluate *MIRO1* expression in primary human brain tumors, we analyzed scRNA-seq data of 24 Grade IV gliomas, 3 Grade III, 7 Grade II, and 5 non-neoplastic controls (seizure patients) from (*22*), comprising a total of 206,532 CD45- cells (Fig. 1C). *MIRO1* was predominantly expressed in tumors, glial cells, and cycling cells and, similar to PD, showed higher expression in glioma samples than in controls (Fig. 1D, S1A). We next compared differentially expressed genes (DEGs) between *MIRO1⁺* and *MIRO1⁻* tumor cells in untreated Grade IV gliomas (n=116,727 cells) and performed gene set enrichment (GSEA). *MIRO1⁺* cells showed upregulation of pathways associated with protein synthesis and oxidative phosphorylation, while downregulated pathways included cell cycle and G2/M checkpoint (Fig. S1B). Interestingly, GSEA of *MIRO1⁺* and *MIRO1⁻* PD DA neurons (n=2,715) showed little overlap with pathways linked to glioma (Fig. S1C). These data indicate that although *MIRO1* upregulation is a shared feature across brain pathologies, MIRO1 likely plays disease-specific roles and engages different transcriptional programs in neurodegeneration versus cancer.

**Figure 1.**
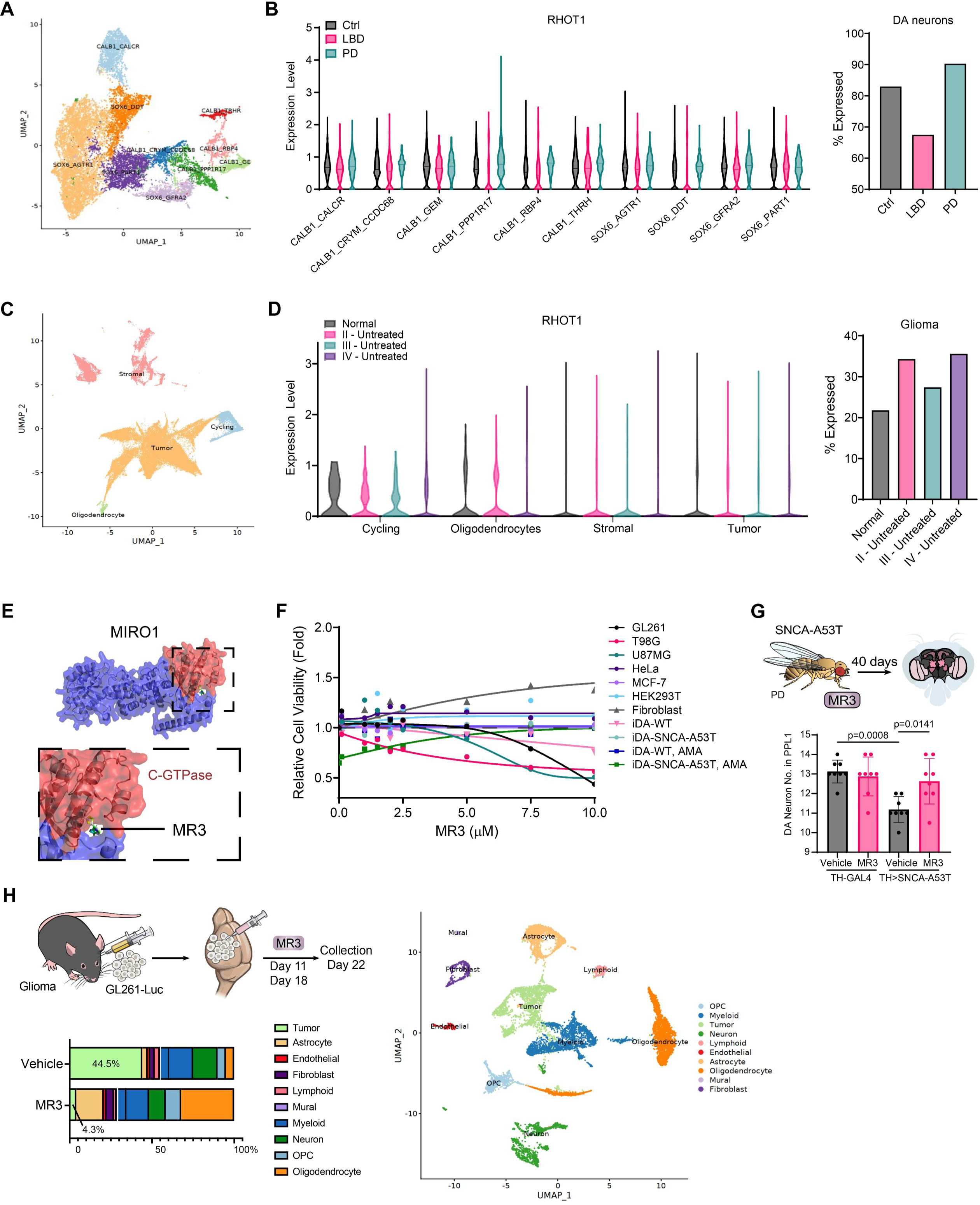
MIRO1 is linked to glioma and PD. **(A)** *MIRO1 (RHOT1)* expression in scRNA-seq datasets of DA neurons from control (Ctrl), LBD, and PD patients (*21*), shown as a UMAP. **(B)** Violin plots showing *MIRO1* expression in each cell cluster across different conditions and bar plots showing the percent of cells expressing *MIRO1* in each condition. **(C)** *MIRO1* expression in scRNA-seq datasets of CD45- cells from glioma and control patients (*22*), shown as a UMAP. **(D)** Violin plots showing *MIRO1* expression in each cell cluster across different conditions and bar plots showing the percent of cells expressing *MIRO1* in each condition. **(E)** Docking of MR3 in the C-GTPase domain of the MIRO1 protein (AF3). **(F)** MTT cell viability assay of glioma (GL261, T98G, and U87MG), non-glioma cells (HEK293T, HeLa, fibroblasts, and MCF7) and iDAs (WT and SNCA-A53T) treated with increasing concentrations of MR3 for 72 hours. For iDAs, AMA (10 μM) was added for the last 6 hours of treatment. Data are presented as the mean fold change in relative cell viability of treated cells vs. vehicle, per cell line (n=6 wells per condition). **(G)** The number of DA neurons in the PPL1 cluster in flies fed with vehicle or MR3 for 40 days. n=8 flies per condition. One-way ANOVA post hoc Tukey test. Error bars represent the mean ± SD. **(H)** Schematic of the GL261 glioma mouse model experimental setup. UMAP plot of all NeuN- nuclei from MR3-treated mice and untreated (vehicle) mice, along with a bar chart illustrating the proportion of each cell type in MR3-treated and -untreated mice.

### A MIRO1-binding compound inhibits glioma cell survival but rescues Parkinsonian neuron loss

The observation that glioma and PD brain tissues exhibit elevated *MIRO1* levels yet display distinct transcriptional programs prompted us to examine the impact of MIRO1 targeting in these disease contexts. We tested the effect of MIRO1 Reducer 3 (MR3), a lead compound from our previously characterized MIRO1-binder series, across a panel of cell lines: three glioma lines – two human (T98G, U87MG) and one mouse (GL261), two non-brain cancer lines (HeLa and MCF-7), and non-cancer controls (HEK293) and healthy donor-derived fibroblasts (Fig. 1E-F) (*18*). Strikingly, MR3 treatment significantly reduced cell viability across all glioma lines in a dose-dependent manner but did not affect non-glioma cancer or non-cancer cells (Fig. 1F). To assess the effects of MR3 on PD, the PD-associated *SNCA-A53T* mutation was introduced into iPSCs from a healthy male donor (WTC11) (*23*) using CRISPR/Cas9-induced homology-directed repair (HDR) (Fig. S2A) (*24*). The mutant line and its isogenic control were then differentiated into highly pure human induced dopaminergic (iDA) neurons via inducible expression of neurogenin-2 (NGN2), followed by directed dopaminergic differentiation, yielding >90% purity (Fig. S2B) (*25*). Acute treatment of WT and *SNCA-A53T* iDA neurons with Antimycin A (AMA), a complex III inhibitor that induces mitochondrial depolarization and increases reactive oxygen species (ROS), key hallmarks of PD pathogenesis, reduced the viability of *SNCA-A53T* neurons but not of isogenic controls (Fig. 1F and S2C) (*26*). Consistent with our prior findings in patient-derived neurons, MR3 prevented AMA-induced cell death in *SNCA-A53T* iDAs (Fig. 1F) (*14, 15, 17–19, 27, 28*). *In vivo*, MR3 also rescued DA neuron loss in a Drosophila PD model expressing human *SNCA-A53T* (Fig. 1G). Conversely, analysis of single-nucleus RNA sequencing (snRNA-seq) data from our previous study (*29*) showed that administering MR3 to a syngeneic mouse glioma model markedly reduced the proportion of tumor cells from 44.5% to 4.3% (Fig. 1H), consistent with its cytotoxic effects in cell models. Together, these data demonstrate that the same MIRO1-binding compound suppresses glioma cell survival and protects PD-relevant DA neurons, suggesting an as-yet-undefined disease-specific role for MIRO1 in regulating cell survival.

### MR3 remodels apoptotic stress signatures *in vivo*

To understand how MIRO1 participates in cell viability pathways, we performed label-free quantitative mass spectrometry (MS) using data-dependent acquisition (DDA) on dissected glioma tumors from mice treated with or without MR3, alongside matched tissue from the corresponding brain region in non-tumor-bearing wild-type (WT) controls (all data are publicly available via a user-friendly browser called MiroScape – https://miroscape.github.io/MiroScape/#/home) (Table S1) (*29*). Proteomic profiling revealed broad remodeling of protein expression in glioma tissue, which was largely reversed by MR3 treatment (Fig. S3A, Table S1). We focused on regulators of cell death and identified apoptosis-related proteins altered in glioma that are restored by MR3. BAX, a pro-apoptotic effector canonically known for its ability to form macropores in the OMM that facilitate cytochrome c release and downstream apoptotic signaling, was elevated, while Mitofusin 2 (MFN2), a key regulator of mitochondrial dynamics known to be downregulated during apoptotic stress, was decreased (Fig. 2A, S3A, Table S1) (*2, 30*). Western blot analysis of the same brain sections used for MS confirmed the alterations in BAX and MFN2 and showed rescue by MR3 (Fig. 2B). In contrast, we did not detect significant MR3-dependent changes in cell death-related proteins using DDA proteomics in iDA neurons derived from *WT* or *SNCA-A53T* iPSCs (https://miroscape.github.io/MiroScape/#/miroProteome/gliomaAndPDCells) (*29, 30*). These findings further support a disease-specific role of MIRO1 in regulating cell death by modulating the expression and/or activity of key OMM factors.

**Figure 2.**
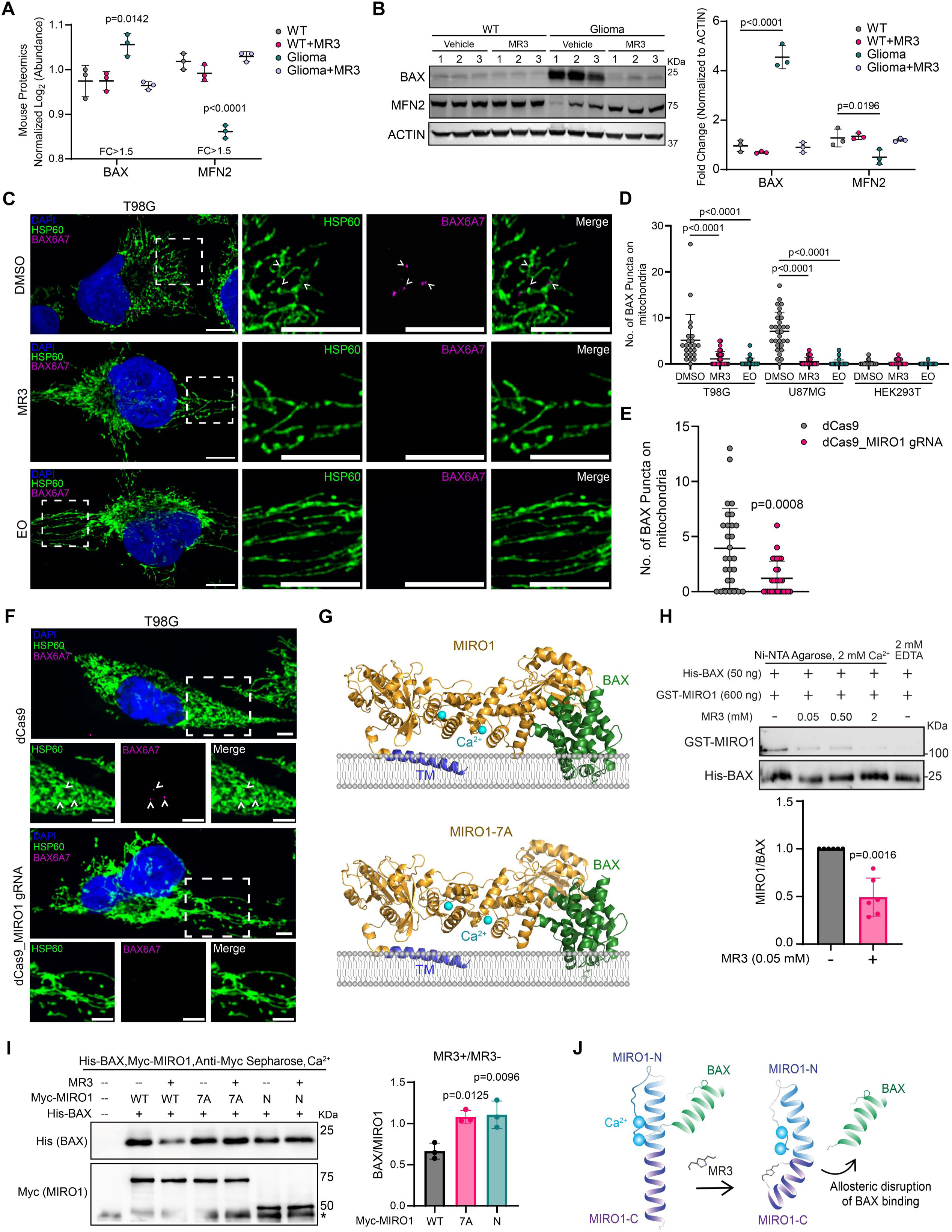
MIRO1 promotes BAX activation. **(A)** Normalized Log_2_ MS spectra abundance of BAX and MFN2 proteins in brain tissues from mice stereotactically implanted with GL261-Luc cells and treated with DMSO (vehicle) or MR3 (10 μM) (Glioma and Glioma+MR3), and from WT mice (not implanted) treated with DMSO or MR3 as shown in Fig. 1H. n=3 mice per condition. One-way ANOVA post hoc Turkey test (compared to WT). **(B)** Representative immunoblots of BAX, MFN2, and ACTIN (loading control) from brain samples as in (A). The band intensity is normalized to ACTIN on the same blot. n=3 mice per condition. One-way ANOVA post hoc Turkey test (compared to WT). **(C)** Representative confocal images of HSP60 and BAX [6A7] in T98G cells treated with vehicle (DMSO), MR3 (10 μM), or EO (10 μM) for 24 hours. Insets show the decrease of the BAX [6A7] signal on mitochondria by MR3 and EO. Scale bars, 10 μm. **(D)** Quantification of the number of BAX [6A7] puncta on mitochondria in T98G, U87MG, and HEK293T cells (non-cancer control) treated as in (C). n=23-30 cells per condition from 3 independent experiments. One-way ANOVA post hoc Dunnett multiple comparisons (compared to DMSO per cell line). **(E)** Quantification of the number of BAX[6A7] puncta on mitochondria from images as in (F). n=28-29 cells per condition from 3 independent experiments. Welch’s unpaired t-test. **(F)** Representative confocal images of HSP60 and BAX [6A7] in dCas9 and dCas9_MIRO1 gRNA T98G cells. Scale bars, 5 μm. **(G)** Modeling of the binding of BAX to WT and mutant (7A) MIRO1 bound to calcium on the mitochondrial membrane. **(H)** *In vitro* binding assay using His-tagged BAX and GST-tagged MIRO1 protein. The MIRO1 band intensity was normalized to His (BAX) IPed on beads to quantify relative MIRO1 binding to BAX in samples treated with vehicle (DMSO) or MR3 (0.05 mM). n=6 independent experiments. Welch’s t-test. **(I)** Lysates from HEK293T cells transfected with Myc-MIRO1-WT (WT), Myc-MIRO1-7A (7A), or Myc-MIRO1-N (N) and treated with DMSO or MR3 (10 μM) for 24 hours were incubated with anti-Myc to IP MIRO1, and co-IP of BAX was assessed by western blot. Asterisk: non-specific bands. The band intensity of BAX was normalized to MIRO1 in each condition. n=3 independent experiments. One-way ANOVA post hoc Turkey test compared to WT. **(J)** Schematic of MR3-mediated allosteric disruption of MIRO1-BAX binding. All error bars represent the mean ± SD.

### MIRO1 is essential for BAX activation in glioma

Based on our previous finding that MR3 altered BAX protein levels in glioma (Fig. 2A-B), we investigated whether MIRO1 influenced BAX activation at the OMM. We used a BAX antibody (6A7) that specifically recognizes the active conformation of monomeric BAX on the OMM, a prerequisite for macropore formation (*2, 31*). BAX can induce either majority or minority mitochondrial outer membrane permeabilization (maMOMP or miMOMP, respectively), defined by the fraction of mitochondria involved (*2*). We found that glioma cells (T98G and U87MG cells) display basal activation of BAX, forming puncta that colocalize with a subset of mitochondria, a hallmark of miMOMP (Fig. 2C-D) (*2*). Treatment with MR3 markedly reduced the number of mitochondria-localized BAX puncta to a level comparable to that of Eltrombopag (EO), a recognized BAX inhibitor (Fig. 2C-D) (33). Knockdown of *MIRO1* in T98G glioma cells using dCas9-CRISPRi with a non-coding-sequence-targeting gRNA (Fig. S3B-C) phenocopied MR3 and EO treatment (Fig. 2E-F), supporting that MIRO1 promotes BAX activation and/or recruitment to the OMM.

### MIRO1 forms a complex with BAX that is allosterically disrupted by MR3

Given that MIRO1 and BAX both localize to the OMM, we hypothesized that MIRO1 directly interacts with BAX to promote its activation. To test this, we immunostained T98G cells expressing Myc-tagged wild-type MIRO1 (Myc-MIRO1-WT) and measured the colocalization of MIRO1 and active BAX puncta on the mitochondrial surface. Treatment with MR3 significantly reduced the number of double-positive puncta (Fig. S3D), indicating that MR3 disrupts the MIRO1-BAX interaction. Computational modeling using AlphaFold3 (AF3) revealed a strong interface between BAX and the N-terminal domain of MIRO1 (amino acids 1-235) (Fig. 2G). *In vitro* binding assays confirmed that MIRO1 binds BAX in a calcium-dependent manner and revealed that MR3 blocks this interaction in a dose-dependent manner (Fig. 2H, S3E). Co-immunoprecipitation (co-IP) assays further confirmed the interaction between BAX and Myc-MIRO1-WT, which was also reduced by MR3 (Fig. 2I). To investigate the nature of this interaction, we mutated the putative MR3-binding residues in MIRO1’s C-terminal GTPase domain to alanine (MIRO1-7A) (*18*). This mutation did not significantly alter the AF3-predicted MIRO1(7A)-BAX interface (Fig. 2G), and MIRO1-7A retained its ability to bind BAX (Fig. 2I). Moreover, expressing an N-terminal fragment of MIRO1 that contains the predicted BAX-binding domain (amino acids 1-395; Myc-MIRO1-N) was sufficient to pull down BAX (Fig. 2I). Neither MIRO1-7A nor MIRO1-N responded to MR3 interference (Fig. 1E, 2I), consistent with the absence of MR3-binding sites in these variants.

These findings show that MIRO1 interacts with BAX through MIRO1’s N-terminal domain and that MR3 allosterically disrupts this interaction by binding to the C-terminal GTPase domain (Fig. 2J), potentially inducing a conformational change that releases BAX.

### MIRO1 mediates mtDNA release via BAX macropores

We next examined the downstream consequences of the MIRO1-BAX interaction. Although BAX macropores are typically associated with apoptosis, glioma cells are proliferative, suggesting that BAX may have non-apoptotic, glioma-specific functions. While maMOMP canonically triggers apoptosis, the miMOMP, as observed in glioma cells, is typically non-lethal (*2*). Intriguingly, MR3 treatment, which suppresses BAX activation, induces glioma cell death (Fig. 1). We hypothesize that miMOMP may be co-opted by glioma cells to promote glioma cell survival. One potential pro-survival mechanism associated with miMOMP is the release of mtDNA into the cytosol, which activates the cGAS-STING pathway, among others, to alter gene expression (*2, 6, 32, 33*). To test whether glioma cells exhibit mtDNA leakage, we performed immunostaining for double-stranded DNA (dsDNA) and mitochondrial markers across several cell types: human glioma cell lines (T98G, U87MG), HEK293T cells, iPSC-induced neurons (iNs) and iDA neurons (*WT* and *SNCA*-*A53T*). At baseline, glioma cells showed more cytosolic DNA foci than non-glioma cells, and this was reduced by MR3 (Fig. 3A-B). Similarly, inhibiting BAX activation with EO reduced cytosolic DNA foci (Fig. 3A-B), linking BAX macropore formation to mtDNA release in glioma. Knocking down *MIRO1* with dCas9-CRISPRi in glioma cells reduced mtDNA leakage (Fig. 3C, S3B-C), further supporting a MIRO1-dependent mechanism. Quantitative PCR (qPCR) analysis of DNA from pure cytosolic fractions of T98G cells confirmed the presence of mtDNA in the cytosol, which was reduced by EO treatment (Fig. S3F).

**Figure 3.**
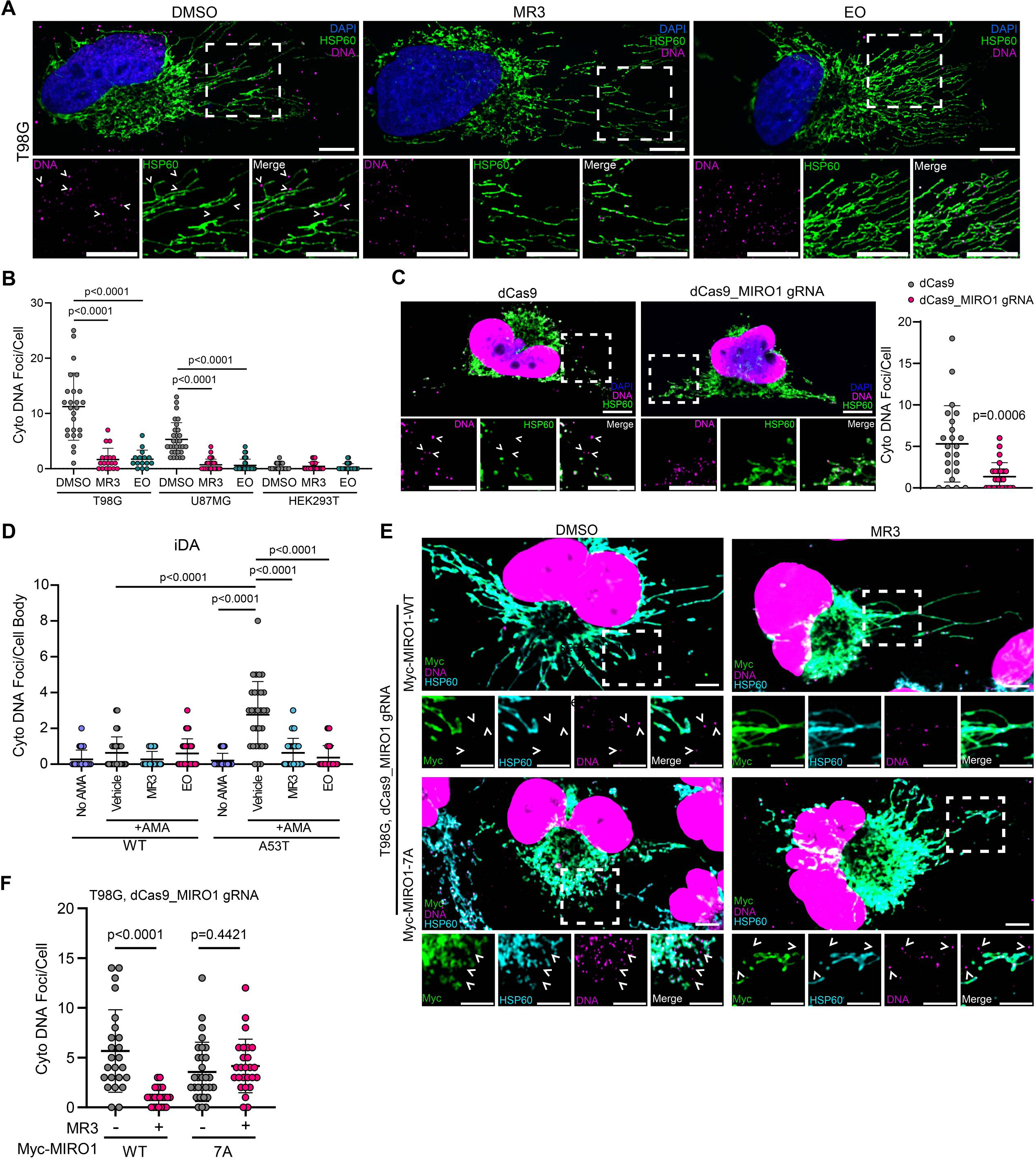
MIRO1 facilitates BAX-mediated mtDNA leak. **(A)** Representative confocal images of dsDNA and HSP60 in T98G cells treated with vehicle (DMSO), MR3 (10 μM), or EO (10 μM) for 24 hours. Insets show cytoplasmic DNA foci. Scale bars, 5 μm. **(B)** Quantification of cytoplasmic (cyto) DNA foci/cell in T98G, U87MG, and HEK293T cells (non-cancer control) treated as in (A). n=15-30 cells from 3 independent experiments. One-way ANOVA post hoc Dunnett multiple comparisons (compared to DMSO per cell line). **(C)** Representative confocal images of HSP60 and dsDNA in dCas9 and dCas9_MIRO1 gRNA T98G cells and quantification of cyto DNA foci/cell as in (B). n=23-25 cells from 3 independent experiments. Welch’s t-test. Scale bars, 10 μm. **(D)** Quantification of cyto DNA foci/cell body in WT and A53T iDAs treated as indicated: Untreated (no AMA), MR3 (10 μM, 30 hours), EO (10 μM, 30 hours), AMA (40 μM, 6 hours). n=30 cells from 2 independent differentiations. Welch’s t-test. **(E)** Representative confocal images of dsDNA, HSP60, and Myc (MIRO1) in vehicle (DMSO) or MR3-treated (10 μM, 30 hours) dCas9_MIRO1 gRNA T98G cells stably expressing either Myc-MIRO1-WT or Myc-MIRO1-7A. Insets show cytoplasmic dsDNA foci. Scale bars, 5 μm **(F)** Quantification of cytoplasmic dsDNA foci/cell from images as in (E). n=23-30 cells from 3 independent experiments. Welch’s t-test. All error bars represent the mean ± SD.

To verify the involvement of MIRO1 in BAX-mediated mtDNA leak beyond glioma cells, we oxidatively stressed iDA neurons with AMA to induce mtDNA leak in *SNCA*-*A53T* neurons (Fig. 3D, S2A-B) (*14, 15, 17–19, 27, 28*). Both EO and MR3 significantly reduced mtDNA leakage, mirroring the involvement of the MIRO1-BAX complex in mtDNA release in glioma cells (Fig. 3D). We also chemically induced BAX macropore formation in iNs with ABT-737/Actinomycin D/qVD-vPh (AAQ), a cocktail that activates BAX and permeabilizes mitochondria while blocking apoptosis (*34*). AAQ-treatment induced significant mtDNA release in iNs, which was markedly reduced by MR3 (Fig. S3G). These data validate the role of the MIRO1-BAX complex in facilitating basal mtDNA leak in glioma cells and stress-induced mtDNA release in neurons.

To determine whether MR3’s effect requires direct MIRO1 binding, we re-expressed either MIRO1-WT or MIRO1-7A in *MIRO1*-deficient T98G cells (Fig. 3E, S3B-C). MR3 reduced mtDNA leak in cells expressing MIRO1-WT, but not in MIRO1-7A-expressing cells (Fig. 3E-F), aligning with our previous findings showing MIRO1-7A maintains BAX binding and is impervious to MR3 (Fig. 2I). Thus, MR3 directly binds MIRO1 to suppress mtDNA leak in glioma. To determine whether MIRO1-mediated mtDNA signaling occurred *in vivo*, we examined activation of the cGAS-STING pathway, a key cytosolic mtDNA-sensing axis, in glioma brain tissues. Western blotting revealed upregulation of key components of the cGAS-STING pathway, which were significantly reduced by MR3 (Fig. 4A). In parallel, MIRO1 protein levels were elevated in glioma tissue and were lowered upon MR3 treatment (Fig. 4A). Crucially, protein measurements by proteomics and immunoblotting showed that MR3 had minimal impact on wild-type, non-tumor brain tissue (Fig. 4A, Table S1). This tissue specificity likely reflects MIRO1 gain-of-function in glioma cells, leading to distinct binding partners or post-translational modifications that MR3 may disrupt. Together, these data position MIRO1 as a central mediator of mtDNA leak via BAX macropores under pathological conditions and highlight the potential of MIRO1 binders as *in vivo* modulators of mtDNA signaling.

**Figure 4.**
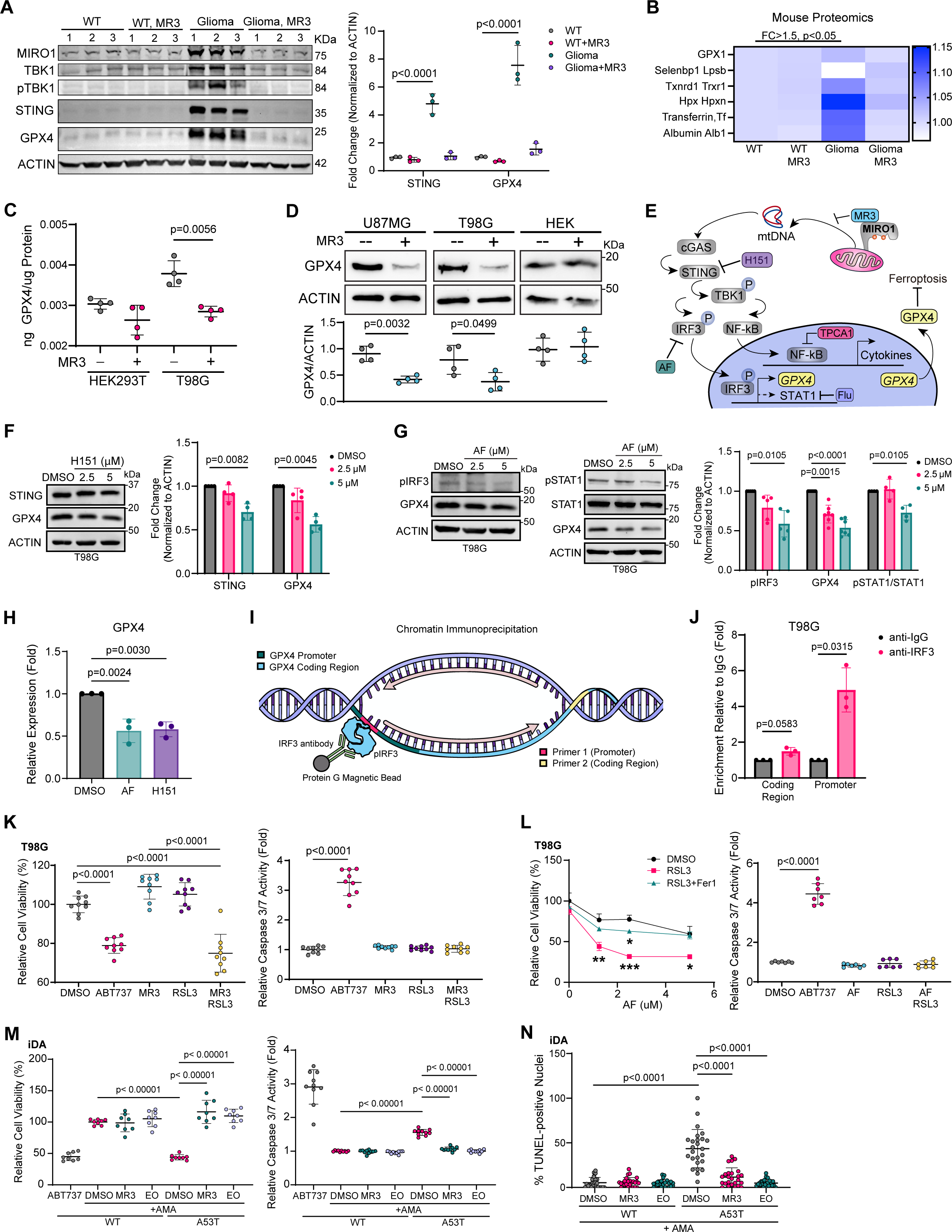
The MIRO1-mtDNA pathway has opposing effects in glioma and PD DA neurons. **(A)** Immunoblotting of MIRO1, TBK1, phospho-TBK1, STING, GPX4, and ACTIN (loading control) in mouse brain samples as in Fig. 2B. STING and GPX4 band intensity were normalized to ACTIN intensity. n=3 mice per condition. One-way ANOVA post hoc Turkey test compared to WT. **(B)** Heat map showing the mean normalized log_2_ MS spectra abundance of selenium and iron metabolism-related proteins from mouse brain tissue, as in Fig. 2A. **(C)** Amount of GPX4 protein (ng/μg of total protein) in HEK293 and T98G +/- MR3 as measured by ELISA. n=4 independent experiments. Welch’s t-test. **(D)** Representative immunoblots of GPX4 and ACTIN (loading control) in U87MG, T98G, and HEK293 cells +/- MR3 (10 μM, 24 hours). GPX4 band intensity was normalized to ACTIN. n=4 independent experiments. Welch’s t-test. **(E)** Schematic of cytoplasmic mtDNA signaling culminating in the upregulation of GPX4 expression and resistance to ferroptosis. Various pharmacological intervention points, including MR3-mediated inhibition of mtDNA release and H151-mediated downstream signaling, are highlighted. **(F)** Representative immunoblots of GPX4, STING, and ACTIN in T98G cells treated as indicated for 24 hours. GPX4 and STING band intensities were normalized to ACTIN intensity from the same blot. n=4 independent experiments. Welch’s t-test. **(G)** Representative immunoblots of GPX4, phospho-IRF3 (pIRF3), STAT1, and phospho-STAT1 (pSTAT1) in T98G treated with AF as indicated for 24 hours. The GPX4 and pIRF3 band intensities were normalized to ACTIN. pSTAT1/STAT1 intensity was normalized to total ACTIN to quantify STAT1 activation. n=5 independent experiments. Welch’s t-test. **(H)** Relative *GPX4* mRNA expression (normalized to *ACTIN*) in T98G cells treated as indicated for 24 hours, as measured by RT-qPCR. Data points represent the average of 3 technical replicates from 3 independent experiments. One-way ANOVA post hoc Dunnett multiple comparisons. **(I-J)** Chromatin isolated from T98G cells was IPed with anti-IgG or anti-IRF3 and captured using Protein G magnetic beads. Following purification, enriched DNA was analyzed by qPCR using primers targeting promoter and non-promoter regions of the *GPX4* gene locus, and enrichment (fold) was quantified relative to IgG. Data points represent the average of 2 technical replicates from 3 biological replicates. Welch’s t-tests. **(K)** Relative cell viability (MTT) and caspase 3/7 activation in T98G cells treated with DMSO (vehicle), ABT737 (10 μM, 24 hours), MR3 (1 μM, 72 hours), RSL3 (10 μM, 24 hours), or co-treated with MR3 and RSL3. n=9 wells from 3 independent experiments. Welch’s t-test. **(L)** Left: Relative cell viability (MTT) (n=4 wells) on T98G cells treated with the indicated doses of AF (5 μM, 18 hours), RSL3 (10 μM, 18 hours), and Fer1 (10 μM, 3 hours pre-treatment followed by 18 hours of co-treatment). Right: Caspase 3/7 activation (n=7 wells) in T98G cells treated with DMSO (vehicle), ABT737 (10 μM, 18 hours), AF (5 μM, 18 hours), RSL3 (10 μM, 18 hours), or co-treated with AF and RSL3. Welch’s t-test. **(M)** Relative cell viability (MTT) (n=8 wells from 2 differentiations) and caspase 3/7 activation (n=10 from 2 differentiations) in WT and A53T iDAs treated with AMA (40 μM, 6 hours) +/- MR3 (10 μM, 30 hours) or EO (10 μM, 24 hours). One-way ANOVA with post hoc Dunnett’s multiple comparisons (compared to A53T DMSO). **(N)** Percent TUNEL-positive nuclei in WT and A53T iDA’s treated with AMA (40 μM, 6 hours) +/- MR3 (10 μM, 30 hours) or EO (10 μM, 24 hours). n=25 images from 2 differentiations. One-way ANOVA with post hoc Dunnett’s multiple comparisons (compared to A53T DMSO). All error bars represent the mean ± SD.

### The MIRO1-STING pathway supports *GPX4* expression in glioma cells, bypassing inflammation

Although cytosolic DNA and basal activation of the cGAS-STING pathway have been reported in glioma, the tumor microenvironment (TME) paradoxically exhibits immunosuppressive properties (*35*).

In many cases, STING signaling is epigenetically silenced in glioma-infiltrating immune cells, rendering them unresponsive to STING agonists (*35*). This discrepancy led us to hypothesize that glioma cells may hijack mtDNA-STING signaling to support their own survival rather than elicit immunogenicity. Our proteomic datasets showed upregulation of selenium- and iron-binding proteins in glioma (Fig. 4B), consistent with enhanced antioxidant defenses and suppression of ferroptosis (*36*). These changes were reversed by MR3 treatment (Fig. 4B). Western blotting revealed that GPX4, a crucial selenium-binding protein and key regulator of ferroptosis resistance, was elevated in mouse glioma tissue and was significantly decreased by MR3 (Fig. 4A). In cell models of glioma (T98G and U87MG), MR3 significantly reduced GPX4 protein levels but had no effect in non-glioma HEK293T cells, as measured by ELISA and Western blotting (Fig. 4C-D). Importantly, treatment of glioma cells with 10 μM MR3 for 24 hours reduced GPX4 levels without affecting MIRO1 protein (Fig. 4C-D, S4A), consistent with our prior reports showing that MR3 destabilizes MIRO1 only at higher doses, longer exposures, or under mitochondrial depolarization (*15, 18, 19*) (*37*). In addition, the same MR3 treatment caused no detectable change in basal oxygen consumption rate (OCR), basal mitophagy, mitochondrial mass, or membrane potential within assay sensitivity (Fig. S4B-G) (*38*). These findings indicate that MR3’s suppression of GPX4 in glioma cells does not result from signaling triggered by mitochondrial dysfunction or loss of MIRO1 protein, but rather from modulation of MIRO1’s function and signaling capabilities.

Since cytosolic mtDNA is a potent activator of the cGAS-STING pathway, we next examined whether this mtDNA-STING axis regulates GPX4 expression in glioma cells, thereby influencing cell survival. We inhibited key nodes of the cGAS-STING pathway using a series of pharmacological inhibitors and measured their effect on GPX4 expression in glioma cells (Fig. 4E). Inhibiting STING downstream of mtDNA sensing by cGAS with the inhibitor H151 (*35, 39*) led to a dose-dependent reduction in GPX4 protein levels (Fig. 4E-F), confirming the involvement of the cGAS-STING pathway in regulating GPX4. We next examined the two main pathways downstream of STING activation: NF-κB and IRF3 transcription pathways (*40*). NF-κB signaling induces the expression of pro-inflammatory cytokines such as IL-6, while IRF3 drives a wave of gene expression dominated by type I interferons (Fig. 4E). These interferons bind to cell receptors, activating the JAK-STAT pathway and phosphorylating STAT1. Phosphorylated STAT1 (pSTAT1) then promotes a secondary wave of interferon-stimulated gene (ISG) expression (Fig. 4E) (*41*).

Inhibiting NF-κB with TPCA-1 (*42*) decreased IL-6 mRNA levels detected by reverse transcription (RT)-qPCR but did not affect GPX4 protein or mRNA levels (Fig. S5A-B), ruling out the involvement of the NF-κB arm of STING signaling. However, inhibition of IRF3 with Auranofin (AF) (*43*) decreased GPX4 protein levels in a dose-dependent manner (Fig. 4E, G), supporting the involvement of the IRF3 arm of STING signaling in regulating GPX4. Notably, at AF doses that reduced GPX4, STAT1 activation and ISG expression remained unaffected (2.5 μM) (Fig. 4G, S5C). Inhibiting STAT1 with Fludarabine (Flu) (*44*) decreased ISG expression but did not reduce GPX4 protein or mRNA levels (Fig. S5D-E). Furthermore, analysis of snRNA-seq data from our mouse glioma model revealed that *GPX4* and *STAT1* were differentially expressed across cell-type subclusters: *GPX4* was highly expressed in tumor clusters, whereas *STAT1* was expressed at lower levels in tumor clusters and at higher levels in myeloid clusters (Fig. 1H, S5F). Gene ontology (GO) analysis of our *in vivo* mouse proteomics dataset further showed that glioma tissue was enriched for pathways associated with translation, macromolecule biosynthesis, and metabolic processes, but no prominent activation of immune pathways was observed (Fig. S5G, Table S1). Together, these data suggest that STING-pIRF3 signaling sustains GPX4 expression in glioma cells independent of NF-κB or STAT1 inflammatory signaling.

Inhibition of STING and IRF3 not only decreased GPX4 protein levels but also significantly reduced *GPX4* mRNA levels without affecting ISG expression (Fig. 4H, S5C), suggesting that the transcription factor IRF3 may directly regulate *GPX4* transcription. We performed chromatin immunoprecipitation (ChIP) using an antibody against IRF3, followed by qPCR with primers spanning putative IRF3-binding elements within the *GPX4* promoter or coding region (non-promoter) (Fig. 4I). IRF3 exhibited significant enrichment at the *GPX4* promoter compared with IgG control (Fig. 4J), suggesting direct occupancy of the *GPX4* promoter region. These data collectively show that mtDNA release in glioma activates pIRF3-dependent upregulation of GPX4, circumventing mtDNA-induced STAT1 inflammatory signaling.

### Expression of the MIRO1-pIRF3-GPX4 pathway is elevated in human glioma

To assess the clinical relevance of the MIRO1-GPX4 axis in human glioma, we analyzed RNA-sequencing data from the TCGA-glioma and GTEx normal brain datasets using GEPIA3. Transcript levels of *GPX4*, *IRF3*, *TBK1*, *STING*, *BAX*, and *MIRO1 (RHOT1)*, but not *MIRO2 (RHOT2)*, a homolog of *MIRO1*, or *TRAK1/2*, encoding binding partners of MIRO1, were all significantly elevated in glioma compared to normal brain tissue (Fig. S6A). Principal component analysis (PCA), treating these genes as a single set, clearly distinguished normal brains from glioma tissues (Fig. S6B). These expression patterns align with our mechanistic data (Fig. 2-4), which show that the MIRO1-mediated cytosolic mtDNA stress pathway is activated in glioma cells and may prime tumor cells for survival.

### MR3 sensitizes glioma cells to ferroptosis and protects Parkinsonian neurons from apoptosis

To determine if blocking the MIRO1-dependent upregulation of GPX4 sensitized glioma cells to ferroptosis, we applied MR3 to glioma cells at a dose (1 μM) that did not cause cell death (Fig. 1F, 4K). Co-treatment with the ferroptosis inducer RSL3 significantly decreased cell viability in T98G cells, as measured by MTT, indicating sensitization to ferroptosis (Fig. 4K). Similarly, loss of cell viability was enhanced in cells treated with AF in combination with RSL3, and this effect was inhibited by the addition of Ferrostatin-1 (Fer1), a ferroptosis inhibitor (Fig. 4L). The observed loss of cell viability in cells treated with RSL3 in combination with MR3 or AF was independent of caspase activation, consistent with the induction of non-apoptotic cell death and supporting that the MIRO1-STING-pIRF3 axis sensitizes glioma cells to ferroptosis (Fig. 4K-L). In contrast, MR3 prevented apoptosis in PD iDA neurons exposed to oxidative stress (AMA), as evidenced by increased cell viability, reduced TUNEL-positive nuclei, and reduced caspase-3/7 activation in *SNCA-A53T* iDAs treated with AMA (Fig. 4M-N). Treatment with EO yielded results similar to those of MR3, implicating BAX in the induction of apoptosis in AMA-treated PD iDAs (Fig. 4M-N). Collectively, our work highlights the context-specific outcome of targeting MIRO1: promoting ferroptosis in glioma cells via the STING-pIRF3-GPX4 axis while preventing MIRO1-BAX-dependent apoptosis in PD neurons (Fig. 5A).

**Figure 5.**
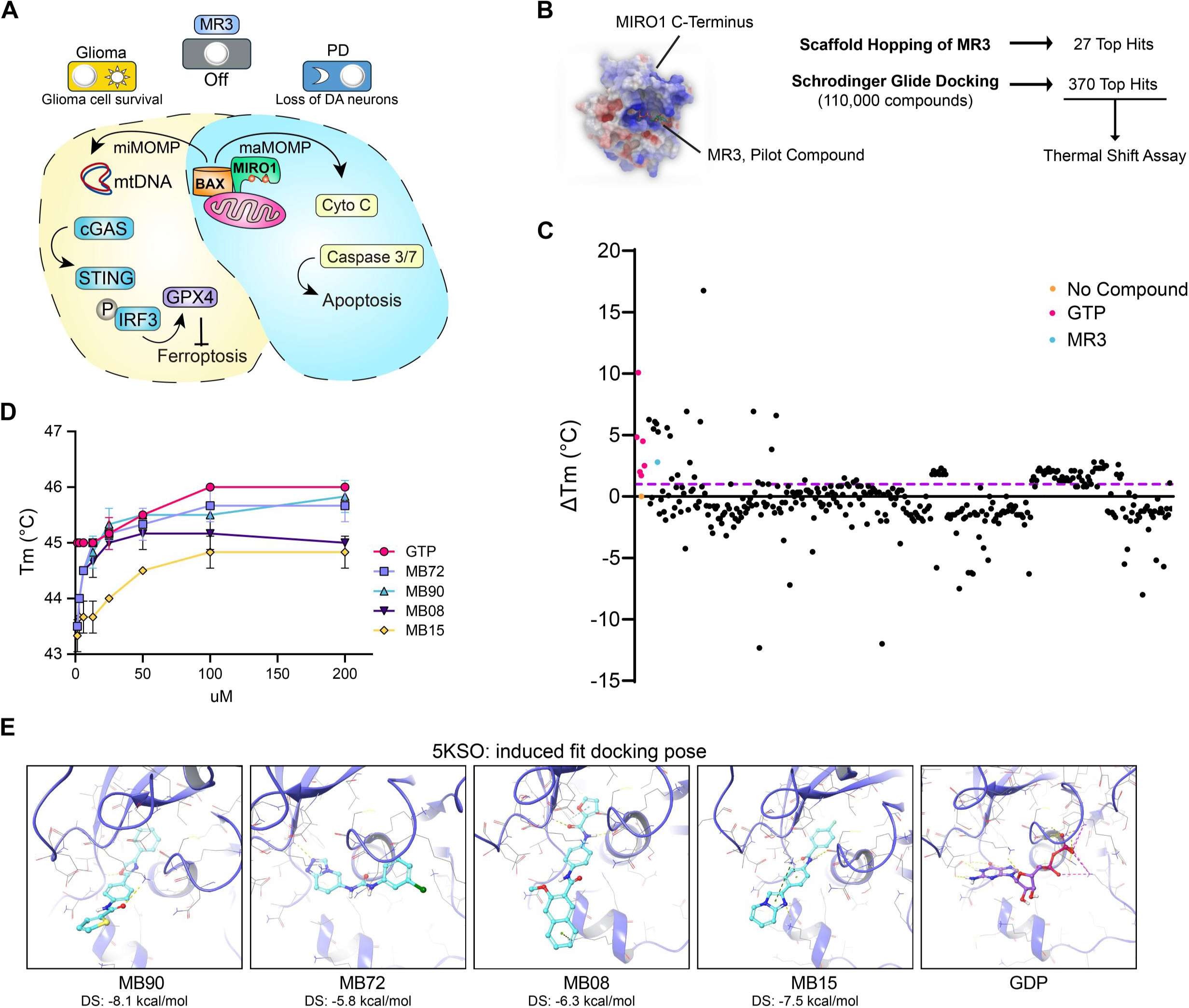
Multiscale drug screens and validations of MIRO1 binders. **(A)** Model of MIRO1-mediated control of cell survival in distinct pathological settings. In glioma, MIRO1 promotes mtDNA release via BAX pores, leading to activation of the STING-IRF3 pathway, upregulation of GPX4, and resistance to ferroptosis. In PD, MIRO1-BAX promotes apoptosis in response to AMA-induced mitochondrial stress. Targeting MIRO1 yields beneficial outcomes across diverse disease contexts, inducing cell death in glioma and promoting survival in PD neurons. **(B)** Schematic of the strategies used to identify novel MIRO1 binders. **(C)** The ΔTm was calculated using TSA for the top 345 compounds identified using the strategies in (B). The ΔTm of GTP controls and MR3 are shown in magenta and purple, respectively. **(D)** TSA was conducted using increasing concentrations of the top 4 compounds. GTP (control) is shown in magenta. **(E)** Induced fit docking of the top 4 compounds and GDP into MIRO1’s C-terminal GTPase domain (PDB: 5KSO) (residues 411-592). All error bars represent the mean ± SD.

### Multi-scale compound screens identify chemically diverse MIRO1-binding analogs

MR3’s selective, allosteric disruption of the MIRO1-BAX interaction and its context-dependent effects in glioma and PD models demonstrate the therapeutic potential and broad applicability of MIRO1-binding compounds (Fig. 5A). We aimed to expand the chemical diversity of MIRO1 ligands to support future therapeutic development in neurodegeneration, brain malignancies, and beyond. We explored structure-activity relationships (SAR) via scaffold hopping and atom/group shape-based similarity screening and performed in silico docking using Schrodinger Glide from a library consisting of >110,000 compounds (Fig. 5B). From these methods, we selected 397 promising candidates based on their performance in the screens, chemical variations, unique binding modes, and distinct predicted protein-ligand interactions (Table S2). These compounds were experimentally evaluated using a thermal shift assay (TSA) against a purified human MIRO1 C-terminal fragment (amino acids 408-592), which included the GTPase domain and exhibited superior TSA stability among several constructs tested (Fig. 5C). Prior work has shown that MIRO1’s C-terminal GTPase activity is not required for TRAK-MIRO1 binding or mitochondrial transport (*45, 46*), highlighting the therapeutic value of targeting this domain without compromising normal mitochondrial function. Of 397 compounds, 345 yielded reproducible thermal shifts (Fig. 5C). Notably, 83 compounds (24.06%) induced a change in melting temperature (ΔTm) ≥1 °C relative to the no compound control, and 141 compounds (40.87%) exhibited a ΔTm of ≤ 1 °C, indicating a robust hit rate (Fig. 5C).

From this pool, we prioritized nine chemically diverse analogs for further validation (Table S2). Eight of the nine compounds displayed a dose-dependent increase (200 μM vs. 1.5 μM) in ΔTm, while the ninth decreased Tm when tested at the higher dose. The top four compounds (ΔTm ≥1.5 °C) showed pronounced, dose-dependent stabilization profiles (Fig. 5D, Table S2), supporting direct engagement with the MIRO1 C-terminal GTPase domain. Rigorous Extra Precision docking calculations revealed docking scores ranging from -5.8 to -8.1 kcal/mol, indicating energetically favorable binding, with predicted binding affinity ranging from moderate to relatively strong (Fig. 5E). Importantly, none of the top four compounds or MR3 inhibited MIRO1’s overall *in vitro* GTPase activity at the therapeutic dose (Fig. S6C-D), consistent with our previous finding (*18*).

We next evaluated the effect of the top four compounds on cell viability in glioma and PD (Fig. 6A). All four analogs showed equal or higher effectiveness than MR3 in decreasing glioma cell viability at all tested concentrations after 24 hours of treatment (Fig. 6A). In *SNCA-A53T* iDA neurons treated with AMA, all four compounds significantly rescued cell viability to a similar or greater extent than MR3 (Fig. 6B). We next tested the *in vivo* pharmacokinetic profiles of two of our four hits: MB72 and MB08. MB72 exhibited an extremely high clearance rate, a short half-life, and was undetectable in the brain (Table S3). By contrast, MB08 demonstrated moderate clearance and tissue distribution and can cross the blood-brain barrier to accumulate in the brain, consistent with a central nervous system (CNS)-active compound (Table S3). We next induced maMOMP in T98G cells using the AAQ drug cocktail to assess MB08’s ability to inhibit BAX activation (*34*). MB08 reduced the BAX6A7 signal on mitochondria to a similar extent as MR3 and EO (Fig. 6C-D). We observed drastic remodeling of mitochondrial membrane structure upon BAX activation in T98G cells, including the formation of donut-like mitochondrial rings detected by confocal microscopy (Fig. 6E) (*34*) and localized OMM discontinuation and evagination into the cytosol (ballooning) visualized by cryogenic electron tomography (Cryo-ET) (Fig. 6F, Movie S1). These restructured membranes are likely caused by BAX complex assembly and pore formation in the OMM. Notably, inhibition of BAX activation with EO, MR3, and MB08 (Fig. 6C-D) significantly reduced the number of donut mitochondria per cell (Fig. 6G), supporting a causal link of MIRO1-mediated BAX activation to mitochondrial membrane remodeling. Together, these findings identify MIRO1 as a regulator of BAX pores and associated membrane destruction and provide a mechanistically grounded set of MIRO1 binders for non-competitive, allosteric disabling of the MIRO1-BAX interface across neurodegenerative and oncogenic contexts.

**Figure 6.**
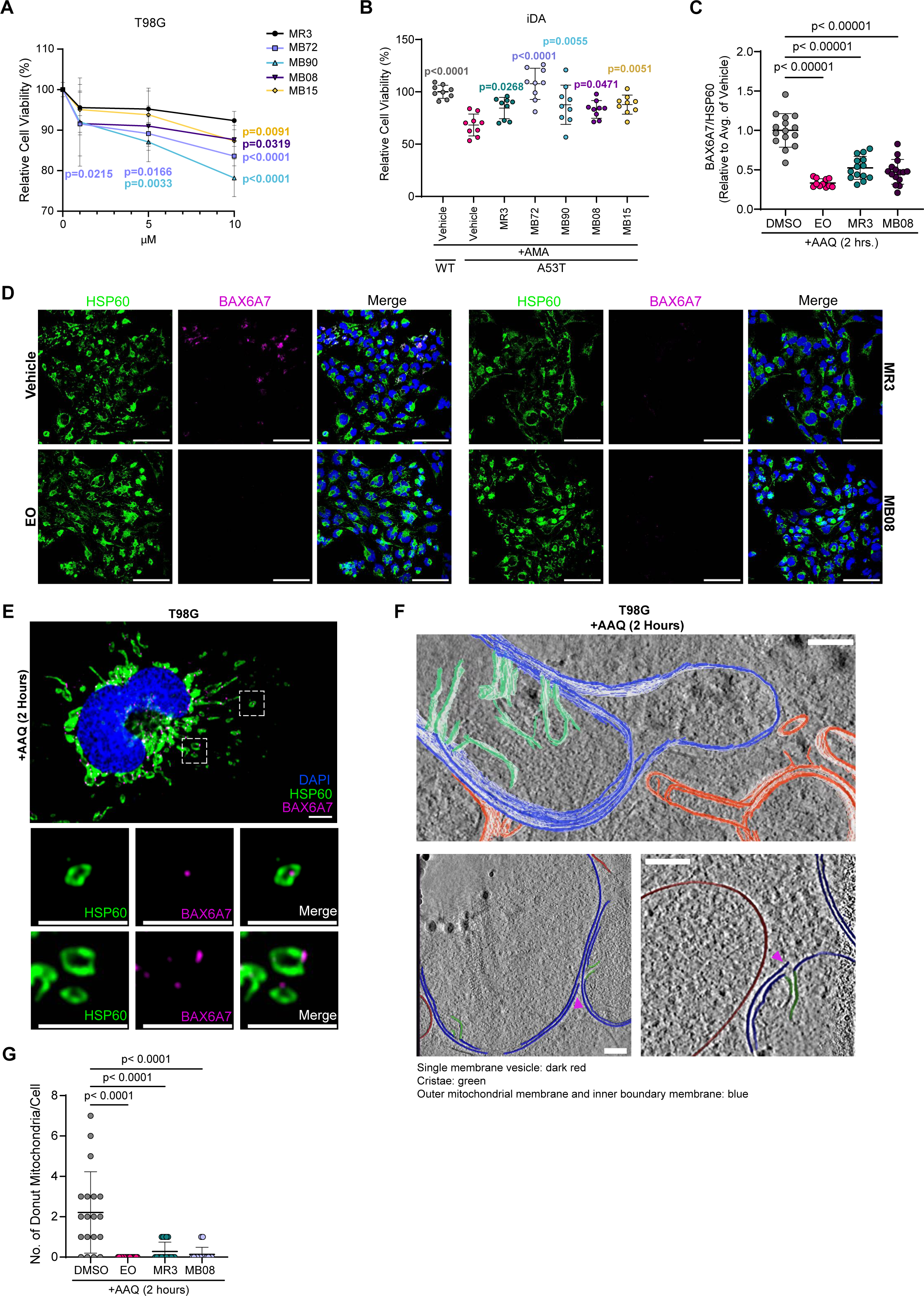
Novel MIRO1 analogs block BAX activation and membrane destruction. **(A)** Relative cell viability (MTT) of T98G cells treated with DMSO or increasing concentrations of the indicated compounds for 24 hours. n=12 wells (DMSO) and n=10 wells (treatments). Multiple unpaired t-tests (compared to MR3). **(B)** Relative cell viability (MTT) of WT and A53T iDAs treated as follows: AMA (40 μM, 6 hours), all other treatments (10 μM, 30 hours). n=9 wells from 2 differentiations. One-way ANOVA with Dunnett’s multiple comparisons (compared to A53T vehicle). **(C)** Relative BAX [6A7] immunofluorescent signal normalized to HSP60 (mitochondria) signal in T98G cells treated with DMSO (vehicle), EO (10 μM), MR3 (10 μM), or MB08 (10 μM) for 24 hours. Cells were co-treated with AAQ (**ABT-737,** 10 μM; **Actinomycin D,** 1 μM**; qVD-vPh,** 10 μM) for 2 hours to induce BAX activation. n=11-15 images from 2 independent experiments. One-way ANOVA with Dunnett’s multiple comparisons (compared to DMSO). **(D)** Representative confocal images of HSP60 and BAX [6A7] treated as in (C). Scale bars, 100 μm. **(E)** Representative confocal images of HSP60 and BAX [6A7] in T98G cells treated as in (C). Insets show donut-like mitochondria (HSP60) and BAX [6A7] puncta. Scale bar, 10 μm (insets: 5 μm). **(F)** Representative Cryo-ET tomograms of T98G cells treated with AAQ for 2 hours as in (C). Scale bars: 100 nm. (**G**) Quantification of the number of donut mitochondria per cell in T98G cells treated with AAQ, as in (C-E). n=15-19 cells per condition from 3 independent experiments. One-way ANOVA with Dunnett’s multiple comparisons (compared to DMSO). All error bars represent the mean ± SD.

## Discussion

We identify a previously uncharacterized function of MIRO1 as a disease-specific gatekeeper of BAX macropore signaling, expanding MIRO1’s molecular skill set (*10–14, 47*). MIRO1 binds BAX via MIRO1’s N-terminal domain to promote macropore formation and mtDNA release, but with opposing outcomes: in glioma, this activates a STING-pIRF3-GPX4 axis that confers ferroptosis resistance while bypassing inflammation; in Parkinsonian neurons, it triggers oxidative stress-induced apoptosis. Our structure-guided chemical toolkit disrupts this interaction, providing a common therapeutic strategy across diseases. Our work provides insights into the molecular logic by which cells exploit and interpret OMM permeabilization. While BAX macropores canonically trigger apoptosis by releasing cytochrome c, we demonstrate that the same structure can drive non-apoptotic survival signaling when gated by MIRO1. The discovery that glioma cells co-opt basal BAX activation (miMOMP) to sustain GPX4 expression may explain how these tumors maintain chronic activity of the cGAS-STING pathway while remaining immunologically silent (*35*). By decoupling IRF3-mediated transcription from STAT1-dependent inflammation, gliomas gain survival benefits while evading immune surveillance.

Cytoplasmic mtDNA triggers a myriad of signaling cascades. Here, we illuminate a previously unreported, context-specific mtDNA signaling modality: transcriptionally activation of GPX4 and resistance to ferroptosis. This cellular principle has immediate therapeutic implications for glioma. Ferroptosis bypasses resistance mechanisms, such as upregulated DNA repair in glioma stem cells, and has the potential to enhance tumor immunogenicity (*48, 49*). Combining MIRO1 binders with ferroptosis inducers may hold promise as a therapeutic intervention for glioma. In the context of neurodegeneration, mtDNA functions as a potent damage-associated molecular pattern that can trigger the spread of PD-like pathology and neuroinflammation and exacerbate neuronal cell death (*50*). Our finding that MIRO1 binders simultaneously block mtDNA release and prevent apoptosis suggests dual therapeutic benefits: suppressing inflammatory cascades that drive neurodegeneration while directly protecting neurons from oxidative death. Release of mtDNA via BAX pores has also been shown to drive cellular senescence via cGAS-STING activation (*2, 51*), raising the possibility that MIRO1 binders could selectively eliminate senescent cells or prevent the induction of senescence to alleviate age-related diseases or unmask immune suppression in the TME.

Our findings open new investigative avenues with implications across distinct disease states: elucidating IRF3-mediated regulation of GPX4, mapping cell-type-specific determinants of MIRO1-BAX signaling using multi-omics approaches, and exploring crosstalk among other cell death and survival pathways downstream of MIRO1-BAX activity. Validation in patient-derived organoids and orthotopic models remains crucial, as does pharmacokinetic optimization. The limited CNS penetration of MB72, despite its strong *in vitro* activity, underscores this need, whereas MB08’s ability to cross the blood-brain barrier offers a promising lead. Determining the structure of the MIRO1-BAX complex will help design advanced inhibitors to regulate mtDNA signaling and improve cellular responses. By revealing how a single macropore structure executes opposing cell fates through disease-specific gating, we establish MIRO1-BAX as a druggable molecular switch that counters glioma cell survival tactics and neurodegeneration.

## Author Contributions

A.G.S., C.S.K., S.A.S., Z.Z., I.C. and A.S.D. designed and performed experiments, analyzed data, and made figures. C.S.K. analyzed mass spec data and generated cell lines. K.C. and M.L. conducted mouse experiments. Z.D. built the MiroScape website and analyzed bulk RNA-seq and snRNA-seq data. B.H.B. analyzed scRNA-seq data. A.V., M.C.B., N.B., and I.K. engineered the iPSC line. I.C. and W.C. performed Cryo-ET and analyzed data. X.W. designed and supervised the project with input from all authors, especially M.C.B. and M.L. A.G.S. and X.W. wrote the paper with assistance from all authors.

## Acknowledgements

We thank L. Li, M. Li, P. Katiyar, S. Bhat, S. Chandra, O. Noor for technical support, R. Ford and J. Kulp (Conifer Point) for drug screens, L-M. Joubert and Y. Xu for assisting in Cryo-ET, G. Li, A. Ting, C. Beinat, and D. Reardon for reagents, the Vincent Coates Foundation Mass Spectrometry Laboratory, Stanford University Mass Spectrometry (SUMS; RRID: SCR_017801) for processing MS samples, the Stanford-SLAC CryoET Specimen Preparation Center (SCSC) for assisting in Cryo-ET, and the following funders: National Institutes of Health (NIH) (R35GM16151901, RO1NS128040, and RO1GM143258, X.W.; P30CA124435 and 1S10OD030473, SUMS; U24GM139166, SCSC), Stanford Wu Tsai Neurosciences Institute (Translate Award, X.W., M.L.; Seed Grant, X.W., M.C.B.), and Stanford School of Medicine Dean’s Office (Postdoctoral Fellowship, I.C.). The content is solely the responsibility of the authors and does not necessarily represent the official views of NIH.

## Materials and Methods

### Cell models and cell culture

No human subjects were used in this study. The Stanford Stem Cell Oversight Committee approved the iPSC work. The following cell lines were obtained from ATCC: HEK293T (Female, CRL-3216), T98G (Male, CRL-1690), U87MG (Male, HTB-14), HeLa (Female, CRM-CCL-2), and MCF7 (Female, HTB-22). GL261 (Male) was a gift from Dr. David Reardon. Healthy donor-derived fibroblasts (Male, ND38530) were purchased under a Material Transfer Agreement (MTA) from National Institute of Neurological Disorders and Stroke (NINDS).

Glioma (U87M, T98G, and GL261) and non-brain cancer cell lines (HeLa and MCF-7) were cultured in Dulbecco’s Modified Eagle Medium (DMEM) (11995073, Thermo Scientific) or Eagle’s Minimum Essential Medium (EMEM) (30-2003, ATCC), respectively, supplemented with 10% heat-inactivated regular fetal bovine serum (FBS) (35-011-CV, Corning). Non-cancer (HEK293T) and healthy donor-derived fibroblasts were maintained in DMEM with 10% FBS and 1% Pen-Strep. Induced pluripotent stem cells (iPSCs) were cultured in mTeSR^TM^1 on Matrigel (354277, Corning) coated cell culture dishes. Cells were maintained in a 37°C, 5% CO_2_ incubator with a humidified atmosphere and split every 4-5 days. We frequently perform cellular morphology checks when passaging the cell lines. We sterilized the tissue culture room and equipment every week and monitored cell lines for mycoplasma using mycoplasma detection kits monthly.

### Cell treatments

The following inhibitors were used at the indicated concentration unless otherwise indicated in figure legends: H151 (HY-112693, MedChemExpress) at 5 μM, Auranofin (HY-B1123, MedChemExpress) at 5 μM, Fludarabine (Flu) (HY-B0069, MedChemExpress) at 200 μM, TPCA-1 (HY-10074, MedChemExpress) at 1 μM, RSL3 (S8155, Selleckchem) at 10 μM, Fer1 (HY-100579, MedChemExpress) at 10 μM, AAQ (ABT737–HY-50907, MedChemExpress at 10 μM, Actinomycin D–A9415, Sigma-Aldrich at 1 μM, QVD-OpH/QVD–HY-12305, MedChemExpress at 10 μM), MB72 (Conifer) at 10 μM, MB90 (Conifer) at 10 μM, MB08 (Conifer) at 10 μM, MB15 (Conifer) at 10 μM, Eltrombopag olamine (EO) (100941, MedKoo Biosciences) at 10 μM, Antimycin A (AMA) (A8674, Sigma-Aldrich) at 40 μM, and MIRO1 Reducer (MR3) (7091, Tocris) at 10 μM.

### Plasmids and transfection

The following plasmids were used: mito-mKeima (*38*) (131626, Addgene), pRK5-Myc-MIRO1-WT (*52*), pRK5-Myc-MIRO1-7A (*18*), and pRK5-Myc-MIRO1-395 (N) (subcloned from Myc-MIRO1-WT). As shown in (*18*), MIRO1-7A is a drug-resistant form by mutating K427, N428, S432, Q446, K454, K528, and D530 (predicated as MR3-MIRO1 interacting sites with in silico modeling) to Alanine in the pRK5-Myc-MIRO1-WT construct, and was validated by the lack of response of MIRO1 protein to MR3 in PD cells. For pTRE3G-Myc-MIRO1-WT and pTRE3G-Myc-MIRO1-7A, MIRO1 gene was PCR amplified from the corresponding pRK5-MIRO1 vector with primers:

Forward: 5’-cacttcctaccctcgtaaatagagctagcgccaccATGgagcagaagctgatctccgagg Reverse: 5’-ttttgtaatccagaggttgattgaattcggccctattatcgctgtttcaataatgctttgtacatagc pTRE3G-Puro backbone vector (253793, Addgene) was digested with NheI and EcoR1. Gibson ligation was performed using in-house 2× Gibson Assembly Master Mix: 100 uL 5×ISO buffer (250 μL 1 M Tris pH 7.5, 12.5 μL 2 M magnesium chloride, 50 μL 10 mM dNTP (N0447S, NEB), 0.125 g PEG-8000 (V3011, Promega), 25 μL 100 mM NAD+ (B9007S, NEB), 25 μL 1 M DTT, add millipore water to 500 μL); 0.4 μL 10 U/μL T5 exonuclease (M0363S, NEB); 6.25 μL 2 U/μL Phusion polymerase (M0530S, NEB); 50 μL 40 U/μL Taq ligase (M0208L, NEB); 3.1 μL 500 μg/mL ET SSB (M2401S, NEB); 215.2 μL millipore water. For each Gibson assembly reaction, an equal volume of 2× Gibson Assembly Master Mix was combined with a DNA mixture containing the digested backbone and insert at an approximate backbone-to-insert molar ratio of 1:3. DNA plasmids were transformed into competent bacterial cells (XL-1 Blue, 200249, Agilent) and sequenced after mini-prep (silica membrane mini spin column, 1910-250, Epoch). For high-efficiency transformation of plasmid DNA products from the Gibson assembly, competent bacterial cells were generated using the Mix & Go competent cell transformation kit (T3001, Zymo) according to the instructions. The competent cells were maintained to ensure high transformation efficiency (about 99% for 1 pg of plasmid DNA). All plasmid DNA constructs were confirmed by whole plasmid DNA sequencing by ELIM Biopharmaceuticals (Hayward, CA).

Transfections were conducted using Lipofectamine 2000 or polyethyleneimine (PEI). For transfection of mito-mKeima using Lipofectamine 2000, 0.5 μg of DNA or 5 μL of Lipofectamine 2000 was diluted in Opti-MEM (31985070, Thermo Fisher) medium to a final volume of 50 μL in two separate tubes. Mixtures were incubated at room temperature for 5 minutes, then combined and incubated for an additional 20 minutes. DNA-lipofectamine mixtures were then added dropwise to cells. For transfection of MIRO1 plasmids using PEI, 1 μg of the indicated DNA was added to 100 μL of serum-free DMEM containing 6 μL of PEI. The DNA mixture was incubated for 20 minutes at room temperature and then added to cells in 1.4 mL of DMEM supplemented with 10% FBS. The media was replaced 24 hours post-transfection.

### Lentivirus generation and cell transduction

HEK293T cells were seeded at a density of 1.0×10^6^ cells per well in a 6-well plate. The next day, cells were cotransfected with the lentiviral packaging plasmids pMDLg/pRRE (12251, Addgene), pRSV-rev (12253, Addgene), and pMD2.G (12259, Addgene) (375 ng, 375 ng, and 250 ng) and plasmids carrying genes of interest (1000 ng) using Lipofectamine 3000 (Thermo Fisher Scientific) according to the manufacturer’s instructions. Viral supernatants were harvested at 24 and 48 hours post-transfection and filtered through a 0.45 µm syringe filter. To promote the precipitation of viral particles, PEG-it Virus Precipitation Solution (SBI Inc., Palo Alto, CA, USA, Cat# LV810A-1) was used according to the manufacturer’s instructions. To transduce target cells with lentiviral particles, 100-500 μL of viral suspensions were added to the cells, and after 24 hours of incubation, the medium was replaced with fresh DMEM. Transduced cells were expanded in culture for 1 week and, when applicable, selected with Puromycin (ant-pr-1, InvivoGen). Non-transduced cells were used to determine Puromycin dose and duration. For cells transduced with pTRE3G-Myc-MIRO1-WT or 7A, Myc-MIRO1 expression was induced by treating cells with 150 ng/mL of Doxycycline (D3447-500MG, Sigma-Aldrich) for 48 hours to induce expression of Myc-MIRO1-WT or 7A before cell treatments. Transduction and expression of the target gene were validated using immunocytochemistry or Western blotting.

### Generation of *MIRO1* KD cells

All plasmid DNA constructs were cloned into the pLV lentiviral vector containing dCas9-KRAB-T2A-EGFP under the UbC promoter and target sgRNA under the U6 promoter (71237, Addgene) using standard Gibson assembly (E2621S, NEB). All constructs were confirmed by sequencing. The complementary primer set for the sgRNA oligos (*MIRO1*: aggcggagctggcgctgtcc), designed using an online tool (https://crispr.dbcls.jp/gRNAcalc/?seq=aggcggagctggcgctgtcccgg), with an overhang region (Tm = 55°C), was subcloned into the target vector with BsmBI. The top and bottom strands of oligos were resuspended to a final concentration of 100 μM and annealed in a thermocycler by using the following parameters: 37 °C for 30 minutes; 95 °C for 5 minutes; ramp down to 25 °C at 5 °C per minute. Next, annealed oligos were diluted to 1:200 with clean water. The digested and cleaned vector with BsmBI and the annealed sgRNA oligos were assembled using the Gibson Assembly Master Mix at 55°C for 45 minutes in a thermocycler, then transformed into competent bacterial cells (DH5alpha, 18258012, Invitrogen) or high efficiency NEB® 5-alpha Competent E. coli (C2987I, NEB), and QIAprep Spin Mini-prep Kit (27104, Qiagen) was used for plasmid amplification. Plasmids were sequenced following amplification.

Forward Primer: ttttaacttgctatttctagctctaaaacGGACAGCGCCAGCTCCGCCTggtgtttcgtcctttccac Reverse Primer: gtggaaaggacgaaacaccAGGCGGAGCTGGCGCTGTCCgttttagagctagaaatagcaagttaaaa HEK293 cells cultured in 6-well plates were transfected with lentiviral packaging plasmids and dCas9-KRAB (1000 ng) or dCas9_MIRO1gRNA (1000 ng), and lentiviral particles were collected, precipitated, and applied to target cells as described in the “Lentivirus generation and cell transduction” section. Transduction and *MIRO1* KD were validated by monitoring GFP expression and Western blot analysis.

### Generation of *SNCA-A53T* iPSC and iPSC maintenance

*SNCA-A53T* mutant iPSCs were created by introducing a two-base substitution at codon 53 through CRISPR/Cas9-mediated homology-directed repair (HDR) in WTC11 iPSCs that express dCas9-KRAB and NGN2 under a doxycycline-inducible system (*23*) (guide sequence: gtggtgcatggtgtggcaac). Single-cell clones were validated through Sanger sequencing of HDR-edited cells and expanded to establish stable mutant iPSC lines. Once established, WT and A53T mutant iPSCs were maintained in mTeSR1 medium (85850, StemCell Tech) and grown on Matrigel-coated plates. Passaging was conducted using Accutase (A6964, Millipore). Rock inhibitor (Y-27632 dihydrochloride) (ab120129, Abcam) was added to freshly seeded cells (10 μg/mL). The medium was replaced with fresh mTeSR1 24 hours after seeding and every 24 hours thereafter.

### iDA preparation and maintenance

Human iPSCs were differentiated into induced dopaminergic neurons (iDAs) using a previously established protocol (*25*). Briefly, iPSCs were plated on Matrigel-coated 6-well plates at a density of 6×10^5^cells/well in mTesr1 medium supplemented with Rock Inhibitor (10 μM). The next day, media was replaced with DMEM/F12 (10565-018, Thermo Fisher) supplemented with 1×N2 (17502-048, Thermo Fisher), 1×B27 (17504-044, Thermo Fisher), 1×non-essential amino acids (11140-050, Thermo Fisher), 10 ng/mL BDNF (450-02, PeproTech), 10 ng/mL GDNF (450-10, PeproTech) and 2 μg/mL Doxycycline (Day 0/1 media). Forty-eight hours later, cells were dissociated with Accutase and seeded on plates or coverslips coated with PDL (30 minutes at room temperature, followed by 3 washes with sterile water and allowed to dry for 2 hours) and Laminin (10 μg/mL added to plates or coverslips following PDL coating; incubated at 37°C for two hours before cell seeding) (23017-015, Thermo Fisher). Cells were seeded in STEMDiff Midbrain differentiation medium (100-0038, StemCell Tech) supplemented with 2 μg/mL of Doxycycline, 1 μg/mL mouse laminin and 200 ng/mL of SHH (78065, StemCell Tech). Full media changes were performed from days *in vitro* (DIV) 3-4, and half media changes were conducted from days 5-8 with supplemented differentiation medium. On DIV 9, the medium was replaced with STEMDiff Midbrain Neuron Maturation medium (100-0041, StemCell Tech) supplemented with 2 μg/mL doxycycline and 1 μg/mL mouse laminin. Starting on DIV 10 and every two days thereafter, half of the medium was replaced with supplemented maturation medium. Cell treatments were conducted on DIV 19, and cells were processed for experiments on DIV 20. Purity of iDA preparation was determined by immunocytochemical analysis, as described in the "Immunocytochemistry and confocal microscopy" section, using antibodies against tyrosine hydroxylase (TH), beta-Tubulin-III (TUJ1) and DAPI and by quantifying the percent of TH-positive neurons.

### iNeuron generation and maintenance

Human iPSCs were differentiated into induced neurons (iNs) using a previously established protocol (*53*). Briefly, iPSCs were plated on Matrigel-coated 6-well plates at a density of 5×10^5^ cells/well in mTesr1 medium supplemented with Rock Inhibitor (10 μM). The next day, media were replaced with DMEM/F12 supplemented with HEPES (11330032, Gibco), 1×N2 (17502-048, Thermo Fisher Scientific), 1×non-essential amino acids (11140-050, Thermo Fisher Scientific), 1×GlutaMAX (35050061, Thermo Fisher), and 2 μg/mL doxycycline. Full media changes were performed at 24 and 48 hours post-seeding. Pre-differentiated iPSCs were then dissociated with Accutase and plated on coverslips coated as described in the *"*iDA preparation and maintenance" section at a density of 5×10^4^ cells/well. Cells were seeded in BrainPhys neuronal medium (05790, StemCell Tech) supplemented with 1×B27 (17504-044, Thermo Fisher), 10 ng/mL BDNF (450-02, PeproTech), 10 ng/mL NT-3 (450-03, PeproTech), and 1 μg/mL mouse laminin (23017-015, Thermo Fisher). Half-media changes were performed every 72 hours.

### Mouse brain preparations for proteomics

Female C57BL/6J mice, aged 6–8 weeks, were obtained from The Jackson Laboratory and housed at the Stanford University Animal Facility under the approved Institutional Animal Care and Use Committee (IACUC) protocol. A total of four treatment groups were established for the proteomic experiments: gliomas, gliomas with treatment, wild-type, and wild-type with treatment. Gliomas were induced by implanting GL261-Luc cells into mice, thereby establishing intracranial tumors. Briefly, mice were anesthetized with ketamine (100 mg/kg) and xylazine (10 mg/kg) via intraperitoneal (i.p.) injection, and topical eye gel was applied to their eyes to lubricate them during the procedure. A small midline incision was made to expose the skull, and a burr hole was drilled over the left striatum, located 2 mm lateral to the sagittal suture and 2 mm posterior to the coronal suture. A total of 1.3×10⁵ GL261-Luc cells (2 µL volume in PBS) were stereotactically implanted at a depth of 3 mm from the cortical surface. Tumor growth was monitored on day 7 post-implantation using an IVIS platform (Lago, Spectral Instrument Imaging) following i.p. injection of 200 µL D-luciferin (150 mg/kg for a 20 g mouse (LUCK, Gold Biotechnology). Mice were assigned to treatment groups (MR3 or vehicle) based on luminescence intensity to ensure similar luminescence between groups. On Days 11 and 18 post-implantation, MR3 was administered intracranially at the tumor implantation site at a dose of 10 μM in DMSO (vehicle; final volume 5 μL). Control mice were treated with 5 μL of DMSO solution. For the non-tumor (wild-type) experimental groups, mice were treated with either 5 µL of DMSO or 5 µL of MR3 (10 µM) on days 11 and 18. The intracranial administration was carried out using the same surgical procedure employed for tumor implantation. All experimental mice were euthanized on day 22, and their whole brains were snap-frozen in liquid nitrogen and stored at -80°C. Only the bottom region of the left hemisphere of the brain, where the tumor was implanted, was used for proteomic analysis. For verification of proteomic results using western blotting, mouse brain tissue adjacent to the proteomic sample was dissociated using a homogenizer. Cell lysates were prepared as described in the "Western blot" section. DDA MS was run at SUMS as described in (*37*). All MS/MS samples were analyzed using the MaxQuant platform (version 2.4.2.0) with Andromeda search engine at 10 ppm precursor ion mass tolerance against the SwissProt and TrEMBL Mus musculus proteome database (68,618 entries, UniProt (http://www.uniprot.org/)). The label-free quantification (LFQ) and Match Between Runs were used with the following search parameters: semi-tryptic and Lys-C digestion; fixed modifications with propionamide on cysteine; dynamic oxidation of methionine; protein N-terminal acetylation; acetylation on lysine; and deamidation on asparagine and glutamine as variable modifications. A false discovery rate (FDR) of less than 1% was obtained for both unique peptides and unique proteins. LFQ intensity values were log_2_-transformed for further analysis, including data normalization, and missing values were imputed using values drawn from a normal distribution centered at the detection limit. All further processing was conducted under the Promor package (https://github.com/caranathunge/promor). Raw data are available on the ProteomeXchange Consortium via the PRIDE (*54*) partner repository with the dataset accession no. PXD079657 and project DOI: 10.6019/PXD079657 as of the date of publication or via Reviewer ID (Username; Password: XY5FwA81KgTh).

### Western blot

Briefly, cells were lysed in western blot lysis buffer (50 mM Tris- pH 7.5, 150 mM NaCl, 1% (v/v) Triton X-100, 0.1% SDS (v/v), 1 mM EDTA, and Protease Inhibitor Cocktail III). Following lysis, samples were centrifuged at 13,000 rpm for 10 minutes at 4°C. Protein concentration was measured using the Pierce™ BCA Protein Assay Kit (23225, Thermo Scientific). Lysates for SDS-PAGE were mixed with 4×Laemmli sample buffer (161–0747, Bio-Rad) containing 5–10% β-mercaptoethanol, and then boiled for 5-10 minutes before being loaded (30-50 μg) into either a precast 10% polyacrylamide gel (4561034, Bio-Rad) or a lab-made gel for Tris-glycine SDS-Polyacrylamide gel electrophoresis with Tris-glycine-SDS running buffer (24.8 mM Tris, 192 mM glycine, 0.1% SDS). After electrophoresis, nitrocellulose membranes (1620115, Bio-Rad) were used in semi-dry transfer with Tris-glycine Transfer Buffer (48 mM Tris, 39 mM glycine, 20% methanol (v/v), pH 9.2). Transferred membranes were first blocked in TBST (TBS with 0.01–0.05% Tween 20) with 5% milk or 4% BSA in TBST for 1hoursat room temperature, and then immunoblotted with the following primary antibodies in TBST with 5% milk or 4% BSA at 4°C overnight (dilution: 1:1000): mouse anti-MIRO1 (WH0055288M1, Sigma-Aldrich-Aldrich; PA5-72835), rabbit anti-STAT1 (9172S, CST), rabbit anti-pSTAT1 (9167T, CST), rabbit anti-beta-Actin (5057S, CST), rabbit anti-BAX (2772S, CST), rabbit anti-STING (13647S, CST), mouse anti-GPX4 (67763-1-Ig, Proteintech), rabbit anti-TBK1 (28397-1-AP, Proteintech), rabbit anti-MYC (16286-1-AP, Proteintech), mouse anti-6x-His (ab18184, Abcam), rabbit anti-pIRF3 (4947T, CST), or rabbit anti-MFN2 (M6319, Millipore). The following day, membranes were incubated with secondary antibodies (1:10,000), including Goat Anti-Rabbit IgG (H+L)-HRP (ab6721, Abcam) and Goat Anti-Mouse IgG (H+L)-HRP Conjugate (1721011, Bio-Rad) for 1 hour at room temperature. West Dura ECL Reagent (34075, GE Healthcare) or SuperSignal West Pico PLUS Chemiluminescent Substrate (34075, Thermo Fisher Scientific) was added to membranes according to the manufacturer’s instructions. Membranes were scanned using a Bio-Rad ChemiDoc XRS system. Membranes were stripped using Restore^TM^ Western Blot Stripping Buffer (21059, Thermo Fisher Scientific) according to the manufacturer’s instructions. Membranes were scanned to verify signal stripping before reprobing. Protein expression was quantified using Fiji software.

### *In vitro* immunoprecipitation

Purified His-BAX protein (50 ng; LS-G132677, Lifespan Biosciences) was added to 20 μL Ni-NTA His-Bind Resin (70666, Millipore Sigma-Aldrich) in 500 μL total volume of IP buffer (50 mM Tris, pH 7.5, 300mM NaCl). Beads were washed 2 times with PBS before adding protein. After the addition of protein, the mixture was incubated for 1.5 hours at 4°C on a rotator. Samples were then centrifuged at 2000 rpm for 1 minute. The supernatant was discarded, and the samples were resuspended in IP buffer containing 2 mM EDTA or 2 mM CaCl_2_. GST-MIRO1 (600 ng) (H00055288-P01, Abnova) was then added to the beads, alongside MR3 at experimental conditions. Due to MR3 precipitating at concentrations above 500 μM, for the 2 mM condition, MR3 was first added to fresh IP buffer to prepare a 5 mM stock, then centrifuged at 13000 rpm for 1 minute. The supernatant was added to the beads, which were then incubated for 1.5 hours at 4°C on a rotator. Samples were then washed three times with IP buffer. After washing, beads were resuspended in Laemmli sample buffer and boiled for 10 minutes before being loaded onto a Tris-glycine SDS-polyacrylamide gel for electrophoresis. Western blot was performed as described in the “Western blot” section.

### Co-immunoprecipitation of MIRO1 and BAX

HEK293 cells were seeded at a density of 3×10^5^ in 6-well plates. When cells reached a confluency of 60-80%, they were transfected with the following plasmids: Myc-MIRO1-WT, Myc-MIRO1-7A, or Myc-MIRO1-395 (N) using PEI as described in the "Plasmids and transfection" section and then treated with 10 μM MR3 for 24 hours. Post-treatment, cells from 3 wells were pooled and lysed in 250 μL of Triton X-100 lysis buffer (300 mM NaCl, 50 mM Tris pH 7.5, 1% Triton X-100, Protease Inhibitor Cocktail III). Cell debris was removed by centrifugation at 13,000 rpm for 10 minutes at 4°C. Then, 60 μL of cell supernatant was collected as ’Input’. The remaining supernatant was incubated with 2 μL of rabbit anti-Myc (16286-1-AP, Proteintech) for 1.5 hours at 4°C on a rotator. Next, lysates were mixed with 60 μL of hydrated protein A–Sepharose beads (101041, Thermo Scientific) for 1 hour at 4°C on a rotator. Samples were then centrifuged at 2000 rpm for 1 minute at 4°C. The supernatant was discarded, and beads were resuspended in IP buffer (300 mM NaCl, 50 mM Tris, pH 7.5, and 2 mM CaCl_2_). Samples were centrifuged and washed twice before the beads were resuspended in IP buffer containing 100 μM MR3 and purified human His-tagged BAX protein (LS-G132677, Lifespan Biosciences), then incubated for 2 hours at 4°C on a rotator. Beads were then washed three times with IP buffer. After washing, beads were resuspended in Laemmli sample buffer and boiled for 10 minutes before being loaded onto a Tris-glycine SDS-Polyacrylamide gel for electrophoresis. Western blot was performed as described in the "Western blot" section.

### Immunocytochemistry and confocal microscopy

Cells were seeded on coverslips at a density of 1-5×10^4^ cells on 24-well plates. The next day, cells were treated as indicated. For cells transduced with pTRE3G-Myc-MIRO1-WT or 7A as described in the “Lentiviral generation and cell transduction” section, Myc-MIRO1 expression was induced by treating cells with 150 ng/mL of Doxycycline for 48 hours before cell treatments. Following treatment, cells were fixed with 4% paraformaldehyde (PFA) for 30 minutes at room temperature. Cells were then permeabilized for 1 hour at room temperature with 0.25% (v/v) Triton X-100 in PBS. Samples were then blocked in PBS containing 3% BSA for 60 minutes at room temperature, followed by incubation with primary antibodies overnight at 4°C. The following antibodies were diluted in PBS containing 1% BSA and 0.1% Triton X-100: rabbit anti-TOM20 (42406S, CST) at 1:200, mouse anti-dsDNA (ab27156, Abcam) at 1:500, Mouse anti-BAX [6A7] (sc-23959, Santa Cruz) at 1:200, Rabbit Anti-Myc (16286-1-AP, Proteintech) at 1:500, Rabbit anti-MIRO1 (HPA010687, Sigma-Aldrich) at 1:200, and chicken anti-HSP60 (CPCA-HSP60, EnCor Biotech) at 1:1000. Samples were then washed in PBS 3 times for 10 minutes/wash with gentle shaking at room temperature before being incubated for 1 hour with the following secondary antibodies diluted (1:1000) in blocking buffer: Alexa fluor 488, Alexa Fluor 568, or Alexa Fluor 647. Samples were washed 3 times for 10 minutes each at room temperature with gentle shaking in PBS. Coverslips were mounted on glass slides using Prolong^TM^ Glass Antifade Mountant with NucBlue^TM^ Stain and left to cure overnight at room temperature.

For iDAs, cells seeded on glass coverslips were fixed with pre-warmed (37°C) 4% PFA for 15 minutes, then washed three times with 1×PBS. Cells were permeabilized with 0.3% Triton-X 100 in PBS for 10 minutes at room temperature and blocked in 5% BSA in PBST (0.1% Triton-X 100 in PBS) for 1 hour. Cells were then immunostained with the following antibodies at the indicated dilutions: chicken anti-HSP60 at 1:1000, mouse anti-dsDNA at 1:200, rabbit anti-MAP2 (CPCA-MAP2, Encor Bio) at 1:10,000, rabbit anti-TH (RPCA-TH, EnCor Biotech) at 1:10,000, and mouse anti-beta-Tubulin III/TUJ1 (T8669, Millipore) at 1:1000. Cells were washed three times for 5 minutes/wash with 1 mM EDTA in PBST and incubated with Alexa Fluor secondary antibodies (1:500) for 45 minutes at room temperature in the dark. Cells were then washed three times with 1×PBS for 5 minutes/wash, and coverslips were mounted on glass slides using Prolong^TM^ Glass Antifade Mountant with NucBlue^TM^ Stain and left to cure overnight at room temperature.

Fixed cell samples were imaged at room temperature with a 63×/N.A. 1.30 oil Plan-Apochromat objective on a Leica SPE laser-scanning confocal microscope (JH Technologies) unless otherwise indicated. Images were processed in Fiji using only linear adjustments to contrast and color. For live-cell imaging of mitochondrial membrane potential, T98G cells were seeded onto coverslips at a density of 1×10^^5^ cells/well in a 24-well plate. The next day, cells were treated with DMSO (vehicle) or 10 μM MR3 for 24 hours. Cells were then incubated with the following dyes for 30 minutes: TMRM (50 nM) and MitoTracker Green (100 μM). Samples were imaged using a 63× water immersion objective at zoom 1-1.5.

### Extraction of RNA from cultured cells and RT-qPCR

Total RNA was extracted from cultured cells using the PureLink RNA Mini Kit (12183018A, Invitrogen) according to the manufacturer’s instructions. Total RNA concentration was measured using a NanoDrop. 1 μg of total RNA was then subjected to reverse transcription using the iScript Reverse Transcription Supermix (1708841, Bio-Rad) according to the manufacturer’s instructions. Before use, cDNA was diluted 1:25 in nuclease-free water. RT-qPCR was performed using the CFX96 Real-Time PCR Detection System (Bio-Rad, Hercules, CA). Each reaction contained the following components: diluted cDNA (4.5 μL), forward primer (10 μM) (0.25 μL), reverse primer (10 μM) (0.25 μL), and Sso Advanced SYBR Green Supermix (1725270, Bio-Rad) (5 μL). The following thermal cycling protocol was used: 95°C for 30 seconds, followed by 35 cycles at 95°C for 15 seconds, and 60°C for 30 seconds. Technical triplicates were assessed for each biological replicate and normalized against the internal standard gene (GAPDH or ACTIN) threshold cycle (Ct) value to calculate ΔCt. The ΔCt of each sample was compared to the ΔCt of the control sample to generate the ΔΔCt value. The relative mRNA expression for each target was then analyzed using the 2^ΔΔCt^ method, and the fold change was plotted relative to control samples.

The following primer pairs were used (ELIM Biopharmaceuticals, Hayward, CA):

RHOT1 forward: 5ʹ- GGTATATGATGTCAGCAATCCCAAATCCT -3ʹ

RHOT1 reverse: 5ʹ- GTATTCTTGTTTAACTTCATGCAGGTCTGACT -3ʹ

GPX4 forward 5’- GAGGCAAGACCGAAGTAAACTAC-3’

GPX4 reverse 5’- CCGAACTGGTTACACGGGAA-3’

BST2 forward 5’- CAGAAGGGCTTTCAGGATGT-3’

BST2 reverse 5’- TTCTCAGTCGCTCCACCTCT-3’

IFI44 forward 5’- GGTGGGCACTAATACAACTGG-3’

IFI44 reverse 5’- CACACAGAATAAACGGCAGGTA-3’

IL6 forward 5’- CCTGAACCTTCCAAAGATGGC-3’

IL6 reverse 5’- TTCACCAGGCAAGTCTCCTCA-3’

ISG15 forward 5’- CTCTGAGCATCCTGGTGAGGAA-3’

ISG15 reverse 5’- AAGGTCAGCCAGAACAGGTCGT-3’

ACTIN forward 5’- ACTGGAACGGTGAAAGGTGAC-3’

ACTIN reverse 5’- AGAGAAGTGGGGTGGCTTTT-3’

GAPDH forward: 5ʹ- GTCTCCTCTGACTTCAACAGCG -3

GAPDH reverse: 5ʹ- ACCACCCTGTTGCTGTAGCCAA -3ʹ

### Mitochondrial fractionation and mtDNA quantification

T98G cells were seeded at a density of 1×10^6^ cells/dish in 15-cm dishes. Twenty-four hours later, cells were treated with 10 µM Eltrombopag (EO) or DMSO for 24 hours. Post-treatment, cells from 2 dishes were pooled for each condition, and samples were normalized to equal cell numbers. Cell pellets were washed with PBS, then resuspended in 1 mL PBS. Cells in suspension (50 µL) were reserved as a whole cell lysate sample, while the remaining sample was reserved for fractionation. Whole cell lysate samples were centrifuged at 1500 rpm for 3 minutes and resuspended in 200 µL 50mM NaOH. The samples were boiled at 95°C for 30 minutes, after which 20 µL 1M pH 8 Tris-HCl was added. The samples were then centrifuged at 17,000 × g for 10 minutes, and the supernatant was collected, diluted to a concentration of 10 ng/µL, and used for qPCR analysis of whole-cell mtDNA copy number. Samples reserved for fractionation were centrifuged at 1500 rpm for 5 minutes. The pellet was resuspended in 1 mL of mitochondrial isolation buffer (200 mM Sucrose, 10 mM Tris-MOPS, and 1 mM Tris/EGTA). The cells were homogenized via 30 strokes with a 2 mL glass homogenizer. Cell debris was then removed by centrifugation at 600 g for 10 minutes twice, followed by centrifugation at 9000 × g for 15 minutes. After the final centrifugation, the pellet was collected as the mitochondrial fraction, and the supernatant was treated as the cytosolic fraction and centrifuged at 17000 × g for 10 minutes. The DNAs in the supernatant were purified using QIAQuick Nucleotide Removal Columns (28306, Qiagen), following the manufacturer’s instructions, and then diluted 1:1 in nuclease-free water and used for qPCR with Sso Advanced SYBR Green Supermix (1725270, Bio-Rad) according to the manufacturer’s instructions. The following thermal cycling protocol was used on a CFX96 Real-Time PCR Detection System (Bio-Rad, Hercules, CA): 95°C for 30 seconds, followed by 35 cycles of 95°C for 15 seconds and 60°C for 30 seconds. The qPCR was performed on DNA samples from the whole cell and cytosolic fractions using the following mtDNA (ND1 and DLoop) and nuclear DNA primers (B2M):

ND1 forward 5’-TCTCACCATCGCTCTTCTACT-3’

ND1 reverse 5’-AGGCTAGAGGTGGCTAGAATAA-3’

DLoop forward 5’-CTATCACCCTATTAACCACTCA-3’

DLoop reverse 5’-TTCGCCTGTAATATTGAACGTA-3’

B2M forward 5’-CTTTCTGGCTGGATTGGTATCT-3’

B2M reverse 5’-CAGAATAGGCTGCTGTTCCTAC-3’

Three technical replicates were performed for each biological sample. For whole-cell mtDNA copy number, mtDNA Ct value was normalized to nuclear B2M Ct value, and the relative mtDNA copy number was analyzed using a 2^−ΔΔCt^ method. Cytoplasmic mtDNA samples were subjected to qPCR as described using the mtDNA primers above and normalized to whole-cell mtDNA copy number.

### Chromatin immunoprecipitation and analysis by qRT-PCR

Chromatin immunoprecipitation (ChIP) was performed using the SimpleChIP® Enzymatic Chromatin IP Kit (Magnetic Beads) (9003, Cell Signaling Technology) with minor modifications. T98G cells were seeded in 15-cm culture dishes and harvested at approximately 90% confluency. Cells were cross-linked in a 15 mL conical tube with 1% PFA in DMEM supplemented with 10% FBS for 10 minutes at room temperature with gentle inversion, every 3 minutes. The reaction was quenched by adding 10× glycine solution to a final concentration of 125 mM, followed by a 5-minute incubation. Fixed cell pellets were stored at -80°C until further processing. For nuclei preparation, cell pellets were resuspended in 1× Buffer A (7006, Cell Signaling Technology) supplemented with dithiothreitol (DTT) (7016, Cell Signaling Technology) and 200× protease inhibitor cocktail (7012, Cell Signaling Technology), then transferred to microcentrifuge tubes. Chromatin was digested with 0.75 µL of micrococcal nuclease (10011, Cell Signaling Technology; 2000 gel units/µL) at 37 °C for 20 minutes, with the tubes inverted every 3 minutes. Following digestion, nuclei were lysed by sonication using a QSonica ultrasonic processor (QSonica, Newtown, CT) equipped with a 422-A titanium microtip probe (1/8-inch diameter) at 75% amplitude for three cycles of 10 seconds on and 30 seconds off. Samples were kept on wet ice between pulses. Cross-linked chromatin was immunoprecipitated overnight at 4°C with rotation using 4 µg of either IRF-3 rabbit monoclonal (4302, Cell Signaling Technology) or normal rabbit IgG (2729, Cell Signaling Technology) antibodies (negative control). Immune complexes were captured using ChIP-grade Protein G magnetic beads (9006, Cell Signaling Technology), washed sequentially with low- and high-salt buffers, and eluted in 1× ChIP elution buffer (7009, Cell Signaling Technology) at 65°C for 30 minutes with gentle vortexing. Cross-links were reversed by incubation with 5 M NaCl (7010, Cell Signaling Technology) and proteinase K (10012, Cell Signaling Technology) at 65°C for 2 hours. DNA was purified using the provided DNA purification columns (10010, Cell Signaling Technology) by centrifugation at 17,000 × g, then eluted in 50 µL of DNA elution buffer (10009, Cell Signaling Technology).

Enrichment of immunoprecipitated DNA was analyzed by qRT-PCR using SimpleChIP Universal qPCR Master Mix (88989, Cell Signaling Technology) according to manufacturer’s instructions on a CFX96 Real-Time PCR Detection System (Bio-Rad, Hercules, CA). PCR cycling conditions were as follows: 95°C for 3 minutes, followed by 40 cycles of 95°C for 15 seconds and 60°C for 60 seconds. To evaluate IRF3 occupancy at the predicted binding site, custom primers (ELIM Biopharmaceuticals, Hayward, CA) flanking the region of interest on the *GPX4* promoter were used (forward: 5’-GGGGACACTTTTCTGCGAGT-3’; reverse: 5’-CAGGCCAGACAACCTGAGAA-3’), corresponding to chr19:1,103,646-1,103,723. To control for nonspecific pull-down, a primer set targeting a *GPX4* non-promoter region in exon 4 was used (Forward: 5’-GTTCCTCATCGACAAGAACGG-3’; Reverse: 5’-TCCCTAGAGAGGACCTACCAG-3’), spanning chr19:1,106,398-1,106,476), Enrichment was calculated using the Percent Input Method, where Percent Input = 2% x 2^(C[T]^ ^2%^ ^Input^ ^Sample–C[T]^ ^IP^ ^Sample)^, and C[T] = C_T_ represents the PCR threshold cycle.

### Seahorse

T98G cells were seeded on XF HS Mini cell culture miniplates at a density of 4.0×10^4^ cells/well and allowed to recover overnight in a 37°C, 5% CO_2_ incubator. After overnight recovery, cells were washed with 200 μL/well of fresh XF DMEM pH 7.4 (Agilent Technologies) and incubated with 200 μL/well fresh XF DMEM supplemented with glucose (10 mM), pyruvate (1 mM), and glutamine (2 mM) in a 37°C, non-CO_2_ incubator for 1 hour. Before starting the assay, the assay medium was removed and replaced with 180 μL/well of fresh medium. The miniplate was then loaded into a Seahorse XF HS Mini Analyzer (Agilent Technologies) with the temperature set at 37°C. Samples were treated with 1.5 μM Oligomycin (complex V inhibitor), 2 μM FCCP and Rotenone/Antimycin A (0.5 μM each) (complex I & III inhibitor).

A total of three measurements were taken after each compound was administered. Data were analyzed using Agilent’s Seahorse Analytics online tool. The data was then directly exported to PRISM.

### MTT and Caspase-GLO assays

For MTT experiments comparing the effects of MR3 on various cell lines, glioma (T98G and U87MG), non-glioma cells (HEK293, HeLa, fibroblasts, and MCF7) and iDAs (WT and A53T) were seeded at a density of 1×10^4^ cells/well in a 96-well plate in the appropriate medium for each cell line (described in "Cell model and cell culture" section). The next day, cells were treated with DMSO (vehicle) or varying concentrations of MR3 (0.1, 1, 1.5, 2.5, 5, 7.5, and 10 μM). The media was replaced after 24 hours of treatment. Forty-eight hours later, MR3 was added for an additional 24 hours. For iDAs, AMA (10 μM) was added for the last 6 hours of treatment. Following the completion of treatment, 3-(4,5-dimethylthiazol-2-yl)-2,5-diphenyltetrazolium bromide (MTT) solubilized in DMSO (5 mg/mL) was added to the cells (10 μL MTT/ 100 μL cell culture medium) and incubated for 3 hours at 37°C to allow for formazan crystal formation. The formazan crystals were then dissolved in 200 μL of DMSO, and absorbance was measured at 570 nm using the TECAN plate reader to assess cell viability.

For MTT experiments assessing the sensitivity of T98G cells to ferroptosis inducers, cells were seeded at a density of 8×10^4^ cells and treated as indicated in the corresponding figure legends. For experiments using AF and H151, T98G cells were seeded at a density of 12×10^3^. Cell viability was assessed as described above. For iDA experiments, 1×10^4^ cells were seeded into 96-well plates prepared as described in the "iDA preparation and maintenance" section. On DIV 19, cells were treated as indicated in the figure legends and cell viability was assessed as described above. For Caspase-GLO experiments, the cell culture medium was replaced with 50 μL of fresh medium following cell treatments. Then, 50 μL of complete Caspase-Glo reagent (G8090, Promega) was added to each well. Plates were incubated at room temperature for 1 hour before luminescence was measured using a plate reader. Luminescence was equivalent to caspase 3/7 activation.

### TUNEL assay

iDAs were seeded on glass coverslips in a 24-well plate at a density of 6×10^4^ per well and treated as indicated in the corresponding figure legends on DIV 20. Following drug treatments, cells were fixed in 4% PFA for 15 minutes at room temperature and permeabilized with 0.25% Triton X-100 in PBS. Samples were then processed using the Alexa Fluor 594 Click-iT^TM^ Plus TUNEL assay kit (C10618, Invitrogen) according to the manufacturer’s instructions. Coverslips were mounted on glass slides using Prolong^TM^ Glass Antifade Mountant with NucBlue^TM^ Stain and left to cure overnight at room temperature. Samples were imaged at room temperature with an ACS APO 20.0×0.60 IMM objective on a Leica SPE laser-scanning confocal microscope (JH Technologies) with a zoom of 1.5. Images were processed in Fiji using only linear adjustments to contrast and color. The number of TUNEL-positive nuclei was manually counted using the Fiji “Cell Counter” plug-in.

### Mitochondrial morphology analysis

Mitochondrial “donuts” were quantified using the MiNA (Mitochondrial Network Analysis) (*55*) plug-in in Fiji. Briefly, the selection tool was used to isolate a single cell in a 2D fluorescent image stained with the mitochondrial marker HSP60 and imaged as described in the “Immunocytochemistry and confocal microscopy” section. The fluorescent signal outside the selection was cleared using the “Clear Outside” function to isolate the signal from single cells. MiNA was launched using the following preprocessing configurations: Unsharp Mask (radius: 2, mask weight: 0.6), Enhanced Local Contrast CLAHE (blocksize: 0, histogram bins: 1, max slope: 0), Median Filter (radius: 1), Otsu thresholding, and Skeletonization. Thresholding was validated by visually inspecting the segmentation and mitochondrial tracing. The identification of mitochondrial donuts was validated by comparing MiNA counts with manual counts.

### GTPase assay

T98G cells (1×10^6^) were lysed in lysis buffer for immunoprecipitation (1% Triton-X 100, 50 mM Tris pH 7.5, 300 mM NaCl, 5 mM EDTA) with protease and phosphatase inhibitor cocktail (78440, Thermo Scientific). Cell lysates were incubated with anti-MIRO1 antibody (1:100) (WH0055288M1, Sigma-Aldrich) for 1 hour at 4°C on a rotator. Lysates were then incubated with 60 μL of hydrated protein-A beads (101041, Thermo Scientific) for 1 hour at 4°C on a rotator. Beads were collected by centrifugation at 3,000 g for 2 minutes. and washed twice with 100 μL of lysis buffer. IPed MIRO1 was eluted from the beads by incubation with 60 μL of 0.2 M glycine (pH 2.5) for 10 minutes, followed by centrifugation at 3,000 g for 2 minutes. Supernatants were neutralized by adding 60 μL of Tris (pH 8). IPed protein was quantified using the bicinchoninic acid protein (BCA) assay (23227, Thermo Scientific). IPed MIRO1 (750 ng) was incubated with Vehicle (DMSO), MR3, MB72 (M726-0007), MB90 (L906-0005), MB08 (BAS0802689), or MB15 (BAS1540983) (10 μM) for 1 hour. GTPase activity was then measured using a GTPase Activity kit (KA1610, Abnova) according to the manufacturer’s instructions. The assay was validated using 0-300 ng of purified MIRO1 (GST-MIRO1) (H00055288-P01, Abnova). Omitting GTP or MIRO1 from the reaction abolished the signal (Pi < 3 mM).

### Compound screening

***Compound library and ligand preparation:*** The Conifer Point Library, containing >110k synthetic small molecules, was strategically designed to maximize chemical diversity while retaining some built-in SAR for analog-by-catalog type testing. The compound library was selected from a variety of vendors and NIH collections, including Asinex, ChemBridge, ChemDiv, Hitfinder, NIH Clinical Collections (NCC) and NIH Diversity 5 (Div5). This library was prepared using the LigPrep application from the Schrödinger suite version 2021-4. The ligands were prepared with ionization states adjusted to pH 7.0 ± 0.5. Tautomeric states were also generated during this preparation stage, and specified chiralities were retained while unspecified chiral centers were varied.

***Virtual protein preparation and grid generation:*** Following preparation of the ligands to be screened, a crystal structure of MIRO1’s C-terminal GTPase was selected for screening from the PDB: 5KSO (*56*). This structure, crystallized with residues 411-592 at a resolution of 2.25 Å, features a hydrolyzed GTP molecule in the active site (GDP + the hydrolyzed phosphate). Utilizing the protein preparation workflow application (*57*), missing hydrogen atoms were incorporated and optimized, solvent molecules were deleted, and proper bond orders were assigned with the force field OPLS4(*58*). Next, the receptor grid generation application was used to define a sampling volume for ligand docking. First, the cleaved phosphate and solvent molecules were removed from the structure, as they may interfere with ligand conformational sampling. The GDP molecule at the binding site was selected as the center of the grid, with a XYZ dimension of 20 Å.

***Glide docking and virtual screening workflow:*** The previously prepared Conifer Point Library was screened using a multi-step virtual screening approach to identify potential MIRO1 binders. To efficiently screen the large library, a cascading precision Glide docking workflow was employed (*59*). This method began with a rapid, low-precision High-Throughput Virtual Screening (HTVS) docking stage to quickly filter out most compounds. The top-scoring compounds from HTVS were then advanced to the Standard Precision (SP) docking simulation, which uses a more detailed scoring function and extensive conformational sampling to refine the binding predictions. From there, the top candidates underwent a rigorous Extra Precision (XP) calculation to further refine their docking scores and poses. The final selection of docked compounds was based on a combination of factors. Compounds were initially filtered to ensure a broad range of chemical functionalities was represented. Promising compounds with diverse, robust chemical functionalities and few liabilities were prioritized. Predicted binding poses were reviewed, with preference given to compounds exhibiting unique binding modes or interacting with distinct residues within the target pocket. This method ensured the final selection of compounds from docking featured a variety of stable chemotypes and predicted binding modes.

### Shape screening

Shape-based screening using the Phase shape screening application within the Schrödinger Suite was employed as a key virtual screening protocol for ligand-based computer-aided drug design (*60*). This method is based on the principle that molecules with similar shapes and electrostatic properties are likely to exhibit comparable binding modes and biological activity (*61*). The screening process involves using a known reference molecule, in this case MR3, to identify and retrieve structurally analogous molecules from the 110k compound library. Low-energy conformers were generated using the ConfGen module (*62*). Both typed pharmacophore and typed atom volume scoring simulations were run. ***TSA***: Streptactin-tagged MIRO1 (residues 408-592) (*37*) was expressed in NiCo21 (DE3) competent *E. coli* (C2529H, NEB). Cells were lysed using B-PER^TM^ Complete bacterial protein extraction reagent (89821, Thermo Fisher). The initial protein purification was conducted with amylose resin. Protein was cut with the HRV3C protease in elution buffer (25-50 mM HEPES, pH 7.5, 100 mM NaCl) and monitored via SDS-PAGE. MIRO1 was further purified using the Strep2 tag on StreptactinXT. Glycerol (10%) was added to the post-streptactin elution buffer to prevent protein precipitation. The truncated MIRO1 protein was then concentrated, and biotin was removed using a 10-kDa molecular-weight cut-off centrifugal filter. For TSA, 0.13-0.37 mg/mL of purified, truncated human MIRO1 protein was incubated with either 100 μM of the test compound or GTP (positive control) in a well consisting of a 10 μL solution containing 10×SYPRO Orange dye. Solutions were dispensed into TempPlate 384-well polypropylene PCR plates (USA Scientific) and sealed with optical film. For the “no compound” control, an equivalent volume of DMSO was substituted for the compound. Fluorescence was monitored using a CFX Opus 384 Real-Time PCR System (Bio-Rad Laboratories) with a thermal ramp from 25°C to 95°C at 0.5°C per step. The FRET scan mode was used for SYPRO Orange detection (Ex: 470–480 nm, Em: 560–580 nm). The results were analyzed using CFX Maestro Software (Bio-Rad) to determine the Tm, defined as the inflection point of the thermal denaturation curve and compared against the apo-protein and GTP-bound protein to identify potential binders. All compound screening was performed in triplicate, so the reported Tm values are averages of the triplicate data. Following this initial TSA screen, nine compounds were selected for further validation using dose-response curves to characterize binding affinity.

### sc/sn-RNA-seq analysis

The acquisition and processing of scRNA-seq data from glioma patients and non-neoplastic controls have been described previously (*22*). Raw and processed data are available at the NCBI’s Gene Expression Omnibus (GEO) accession number GSE278456. In addition to the methods described previously, predicted doublets were identified and subsequently removed from the dataset using the DoubletFinder software (*63*). Furthermore, to confirm the identity of tumor cells in the dataset, we used the copy number prediction software SCEVAN (*64*). Pre-processed scRNA-seq data from dopaminergic neurons in control, LDB, and PD patients were downloaded from the GEO database under accession GSE178265 (*21*). To perform differential expression analysis (DEA) on *MIRO1*-positive versus *MIRO1*-negative cells in each cell cluster, we used the Model-based Analysis of Single-cell Transcriptomics (MAST) framework for DE testing (*65*), considering genes as differentially expressed if they had an average log_2_FC of > 0.5 or <- 0.5 and an adjusted p-value of < 0.05. To perform gene set enrichment analysis (GSEA), the fgsea package (*66*) was used. Genes were ranked by their average log_2_FC values identified by DEA. The Hallmark and KEGG gene set databases were utilized for this analysis. Mouse snRNA-seq data were downloaded from GEO database under accession GSE324504 and analyzed as previously described (*29*).

### Cryo-ET

#### Sample preparation and vitrification

Quantifoil 200 mesh R 2/2 grids with gold bars were placed in the center of 35-mm cell culture dishes, UV-sterilized for 15 minutes, and coated with Matrigel (354277, Corning) according to the manufacturer’s instructions. T98G cells were seeded at a density of 3×10^4^ per well containing 2-3 grids. The next day, cells were treated with AAQ for 2 hours. Grids were vitrified by plunge-freezing into liquid nitrogen-cooled liquid ethane using a Leica EM GP plunge freezer (Leica Microsystems) with backside blotting for 8 seconds.

#### Cryo-FIB milling

Vitrified grids were clipped and loaded into an Aquilos 2 cryo-FIB/SEM (Thermo Fisher Scientific) at SCSC for focused ion beam (FIB) milling. Lamellae were milled to a target thickness of approximately 150 nm.

#### Data collection

Vitrified grid specimens were imaged using a Titan Krios transmission electron microscope (Thermo Fisher Scientific) operated at 300 kV, equipped with a Gatan K3 direct electron detector and a BioQuantum energy filter operated with a 20 eV slit width. Tilt series were collected at a nominal magnification of 26,000×, yielding a calibrated pixel size of 3.44 Å/pixel. Acquisition was performed using a dose-symmetric tilt scheme starting at 0° and tilting from −20° to +20° with a 2° angular increment (21 tilts total). The electron dose per tilt was 5 e⁻/Å², resulting in a cumulative dose of approximately 105 e⁻/Å² per tilt series. Data collection was controlled using Tomo5 (Thermo Fisher Scientific) with a target defocus range of −3.0 to −6.0 μm.

#### Tomogram reconstruction and 3D segmentation

Motion correction, Contrast Transfer Function (CTF) estimation, alignment, and 3D tomogram reconstruction were performed using AreTomo3 (*67*). Tomograms were reconstructed using Weighted Back-Projection (WBP) with a binning factor of 4 (binned pixel size 13.76 Å/pixel). For 3D visualization of mitochondrial membrane remodeling, subcellular structures were segmented using MINT (Multimodal Imaging Neural Toolkit; in preparation). All segmentations were reviewed and refined manually within MINT.

### TCGA and GTEx RNA-seq data analysis

Gene expression data were analyzed using GEPIA3 (https://gepia3.bioinfoliu.com), based on RNA-seq data from the TCGA PANCAN (release 2016-09-01) (*68*) and GTEx (release 2016-04-19) datasets (*69, 70*), integrated via the UCSC Xena project (http://xena.ucsc.edu). Expression differences were visualized using log_2_(TPM + 1) values. Statistical significance was assessed using the built-in GEPIA3 differential expression analysis, applying a two-sample t-test. PCA was performed using GEPIA3’s dimensionality reduction module on the selected gene set (*71*).

### Mitophagy measurements

Cells were seeded in black, clear-bottom 96-well plates (3340, Corning) at a density of 1×10^4^ cells/well in DMEM supplemented with 10% FBS. Twenty-four hours post-seeding, cells were transfected using PEI as described in the "Plasmids and transfection" section with the following modifications: 0.125 μg of mito-mKeima was added to 12.5 μL of serum-free DMEM containing 0.373 μL of PEI. The mixture was incubated at room temperature for 20 minutes and then added to wells containing 87 μL of DMEM + 10% FBS. Twenty-four hours post-transfection, the media were replaced with supplemented DMEM containing DMSO (vehicle) or 10 μM MR3. Following 24 hours of treatment, fluorescence was read using a TECAN Infinite M200 multi-detection plate reader. The plate was read using excitation wavelengths between 400 and 594 nm (step size 2nm) and an emission wavelength of 620 nm. Fluorescence at 400 and 586 nm was recorded, and the fluorescence ratio (586/400 nm) was calculated and normalized to vehicle controls.

### Fly experiments

The following fly stocks were used: *TH-GAL4* (Bloomington Drosophila Stock Center), *UAS-SNCA-A53T* (*72*). All fly stocks were maintained as previously described in (*15*). Flies were fed MR3 (2.5 μM) for 40 days as described in (*15, 19*). Adult fly brains were dissected in PBST (0.3% Triton X-100 in PBS) and incubated with fixative solution (4% formaldehyde in PBST) for 15 minutes at room temperature. Fixed samples were incubated in blocking solution (10% BSA in PBST) for 1 h, then immunostained with rabbit anti-TH (AB-152, EMD Millipore) at 1:200 in blocking solution for 36 hours at 4°C on a rotator. After washing, Alexa Fluor 488 (1:1000) was added to the blocking solution, and the mixture was incubated for 1 hour at room temperature in the dark. The DA neuron number was counted throughout the z-stack images of each brain.

### Browser building

All data shown in this study are publicly accessible through our lab’s interactive platform, MiroScape (https://miroscape.github.io/MiroScape/#/home), which integrates multi-level datasets including transcriptomes, mitochondrial surface proteomes, and whole proteomes. The datasets in this paper are located under the “MiroProteome” → “Glioma Mice” tab in MiroScape. Users can search for proteins of interest by gene name and interactively visualize their patterns. MiroScape was built using the React framework for the front-end architecture and react-plotly.js for data visualization. The entire MiroScape website is hosted on GitHub Pages for public accessibility. All source code is openly available at: https://github.com/miroscape/MiroScape.git. Software and algorithms used: React version 9.2.0 (Meta; https://react.dev/), Ant Design version 5.27.4 (Ant Group; https://ant.design/), react-router-dom version 7.9.4 (Remix; https://reactrouter.com/), iframe-resizer version 5.1.5 (Bradshaw; https://github.com/davidjbradshaw/iframe-resizer), react-plotly.js version 2.6.0 (Plotly; https://plotly.com/javascript/).

### *In vivo* pharmacokinetic study

The study was conducted at AAALAC accredited facility of Sai Life Sciences Limited, Hyderabad, India, in accordance with the Study Protocol SAIDMPK/PKM-25-09-1393. All procedures in the present study was in accordance with the guidelines issued by the Committee for the Control and Supervision of Experiments on Animals (CCSEA) as published in The Gazette of India on December 15, 1998. Prior approval of the Institutional Animal Ethics Committee (IAEC) was obtained before initiation of the study.

Briefly, 8 to 12-week-old male C57BL/6 mice (n=24 per compound) were intravenously administered either MB08 or MB72 (vehicle: 5% DMSO, 5% Solutol HS-15, and 90% Saline) at a dose of 1 mg/kg. Approximately 60 μL of blood samples were collected under light isoflurane anesthesia (Surgivet®) from the retro-orbital plexus of a set of three mice at 0.083, 0.25, 0.5, 1, 2, 4, 8 and 24 hours post-dose in pre-labeled tubes containing 20% v/v K2-EDTA as an anticoagulant. Immediately after blood collection, plasma was harvested by centrifugation at 10000 rpm for 5 min at 4°C, and samples were stored at - 70±10 °C until bioanalysis. Following blood collection, animals were anesthetized, and the whole body was perfused transcardially with 10 mL of chilled PBS. Brain and Heart samples were collected from a set of three mice at pre-determined time points. After isolation, tissue samples were rinsed three times in ice-cold PBS (for 5-10 seconds/rinse using ∼5 mL chilled PBS in a disposable petri dish), dried on blotting paper and weighed. The brain and heart samples were diluted (1 part brain: 2 parts ice-cold PBS and 1 part heart: 4 parts ice-cold PBS), homogenized, and stored at -70±10 °C until bioanalysis. The received concentrations (ng/mL) were corrected for dilution factors (3× for brain and 5× for heart), and the final reported concentrations of the indicated compound in each tissue were expressed in ng/g. analyzed with a fit-for-purpose LC-MS/MS method (LLOQ: 1.00 ng/mL for plasma & heart and 2.00 ng/mL for brain). The pharmacokinetic parameters were calculated using the non-compartmental analysis tool of Phoenix® WinNonlin software (Ver 8.6).

### Quantification and Statistical Analysis

Data is expressed as box whisker, interleaved scatter, or interleaved scatter with bars, unless otherwise stated in figure legends. One-way ANOVA was used to compare multiple groups, and Welch’s t-test was performed when comparing two groups, as described in the figure legends. Statistical analysis was performed using Prism software. Schematics were made using Adobe Illustrator. Sample size and experimental replication are described in the corresponding figure legends. No statistical methods were used to predetermine sample sizes, but the experiments and biological replicates were determined based on the nature of the experiments, degree of variations and published papers describing similar experiments. Precise p-values are labeled in figures.

