## Supplementary Figures, Table, Legends for "The MIRO1-BAX Complex Dictates Life and Death at the Mitochondrial Gate"

### Supplementary Figures, Tables, and Movie

### Supplementary Figures

**Figure S1. Analysis of scRNA-seq data.**

**(A)** (Left) Expression of tumor, stromal, oligodendrocyte, and cycling cell markers and (Right) SCEVAN CNV predictions in CD45- cells from glioma and control patients. **(B)** Differential expression analysis (DEA) and gene set enrichment analysis (GSEA) between *MIRO1*-positive (*MIRO1*+) and *MIRO1*-negative (*MIRO1*-) cells in each of the tumor cell clusters. Cell clusters are shown on the x-axis, and genes and gene sets are shown on the y-axis. For each cluster, genes/gene sets up-regulated in *MIRO1*+ cells are shown in red, and genes/gene sets down-regulated in *MIRO1*+ cells are shown in blue. **(C)** DEA and GSEA between *MIRO1*+ and *MIRO*1- cells in each DA neuron cluster in PD patients.

**Figure S2. Generation of the *SNCA-A53T* iPSCs and validation of their induced DA neurons.**

**(A)** Illustration of the target *SNCA* gene segment that includes annotations involved in the CRISPR/Cas9-induced HDR edit. The A53T mutation occurs at position 53 in exon 3 of the *SNCA* gene (red box indicating the GCA-to-ACC edit) and results in a threonine-to-alanine replacement. Location of symmetric and asymmetric HDR donor templates in yellow and blue, respectively, and gRNA in green. **(B)** Representative confocal images of tyrosine hydroxylase (TH) and TUJ1 in WT and SNCA-A53T iDAs. The percentage of TH-positive neurons is shown. n=30 images from 2 differentiations. Scale bars, 20 μm. **(C)** Quantification of mitochondrial membrane potential (TMRM intensity normalized to MitoTracker Green–MTG–intensity) in WT and SNCA-A53T iDA's treated with DMSO (vehicle) or Antimycin A (AMA) (40 μM) for 4 hours. n=25 images from 2 differentiations. One-way ANOVA with Dunnett's multiple comparisons (compared to WT DMSO). All error bars represent the mean ± SD.

**Figure S3. MIRO1's link to BAX activation.**

**(A)** Scatter plot showing normalized expression levels of proteins altered by MR3 in glioma brain tissues analyzed by MS. n=3. **(B)** Relative *RHOT1* mRNA expression (normalized to *GAPDH*) in dCas9 and dCas9_MIRO1 gRNA T98G cells as measured by RT-qPCR. n=3 independent experiments. Welch's t-test. **(C)** Representative immunoblots of MIRO1 and ACTIN (loading control) in dCas9 and dCas9_MIRO1 gRNA T98G cells. MIRO1 band intensity was normalized to ACTIN intensity from the same blot. n=3 independent experiments. Welch's t-test. **(D)** Quantification and representative confocal images of Myc and BAX[6A7] in T98G cells stably expressing Myc-tagged MIRO1 treated with vehicle (DMSO) or MR3 (10 μM) for 24 hours. Insets show BAX[6A7] puncta on mitochondria. n=28-33 cells per condition. Welch's t-test. Scale bars, 10 μm (Insets: 5 μm). **(E)** Negative controls for Fig. 2H *in vitro* IP. **(F)** Relative cytoplasmic mtDNA abundance (*Dloop* and *ND1*) in cytoplasmic fractions from T98G cells treated with DMSO (vehicle) or EO (10 μM) for 24 hours, as measured by RT-qPCR. n=4-5 independent experiments. Welch's t-test. **(G)** Quantification of cytoplasmic DNA foci/cell body in WT iNeurons treated with AAQ (**ABT-737,** 10 μM; **Actinomycin D,** 1 μM**; qVD-vPh,** 10 μM) for 24 hours. n=30-32 cells per condition from 2 differentiations. One-way ANOVA with Dunnett's multiple comparisons. All error bars represent the mean ± SD.

**Figure S4. MR3 does not affect MIRO1 protein levels or mitochondrial functions.**
**(A)** Amount of MIRO1 protein (ng/µg of total protein) in U87MG and T98G cells +/- MR3 as measured by ELISA. n=4 independent experiments. **(B)** OCR profiles measured by Seahorse of DMSO and MR3-treated T98G cells. n=3 independent experiments. **(C)** Relative basal respiration (OCR) from (B). **(D)** Quantification of mitophagy measured by mito-mKeima (mt-mKeima) in T98G, U87MG, and HEK293 cells treated with DMSO (vehicle) or MR3 (10 µM, 24 hours). n=6. **(E-F)** Quantification of mitochondrial mass (intensity of MTG) (E) and mitochondrial membrane potential (TMRM intensity normalized to MTG intensity) (F) in T98G cells treated with vehicle (DMSO) or MR3 (10 µM) for 24 hours. n=21. **(G)** Representative confocal images of T98G cells stained with TMRM and MTG, treated as in (E-F). Scale bars, 10 µm. All error bars represent the mean ± SD.

**Figure S5. The GPX4 pathway is independent of inflammation.**
**(A)** Relative *IL6* and *GPX4* mRNA expression (normalized to *ACTIN*) in T98G cells treated as indicated for 24 hours, as measured by RT-qPCR. Data points represent the average of 3 technical replicates from 3 biological replicates. Welch's t-test. **(B)** Representative immunoblots of GPX4 and ACTIN in T98G cells treated as in (A). GPX4 band intensity was normalized to the ACTIN intensity from the same blot. n=3 independent experiments. **(C)** Relative mRNA expression of the interferon-stimulated genes *BST2*, *IFI44*, and *ISG15* (normalized to *ACTIN*) in T98G cells treated as indicated for 24 hours, as measured by RT-qPCR. Data points represent the average of 3 technical replicates from 3 biological replicates. **(D)** Representative immunoblots of GPX4, phospho-STAT1 (pSTAT1), STAT1, and ACTIN (loading control) in T98G cells treated with DMSO (vehicle) or 200 µM Flu for 24 hours. Band intensities were normalized to the ACTIN intensity. The intensity of pSTAT1 was normalized to total STAT1 to quantify STAT1 activation. n=3 independent experiments. Welch's t-test. **(E)** Relative mRNA expression of *GPX4* and the interferon-stimulated genes *ISG15* and *STAT1* (normalized to *ACTIN*) in T98G cells treated with DMSO (vehicle) or Flu (200 µM) for 24 hours, as measured by RT-qPCR. Data points represent the average of 3 technical replicates from 4 biological replicates. **(F)** Dotplot of *GPX4* and *STAT1* expression in the indicated cluster from Fig 1H. **(G)** Gene ontology analysis (ShinyGO) and expression of proteins associated with translation, metabolism, and hallmarks of tumor growth from proteomics conducted on mouse brain tissues treated as indicated and as described in Fig. 2A. All error bars represent the mean ± SD.

**Figure S6. Human glioma data analysis and GTPase activity of the compounds.**
**(A)** Differential expression of genes involved in mitochondrial transport and stress response between glioma and normal cortex tissues. *TRAK2*, *TRAK1*, *MIRO2 (RHOT2)*, *MIRO1 (RHOT1)*, *BAX*, *STING (STING1)*, *TBK1*, *IRF3*, and *GPX4* were analyzed using GEPIA3. Red and green

lines represent Glioma (n=166) and Normal Brain Cortex (n=110: GTEx cortex n=105 and TCGA peritumor n=5), respectively. Two-sample t-test. **(B)** Principal component analysis (PCA) of glioma and normal cortex samples based on the expression of *MIRO1 (RHOT1)*, *BAX*, *STING1*, *TBK1*, *IRF3*, and *GPX4*. Colors represent sample origins: Brain Cortex (GTEx), Glioma_Tumor, and Glioma_Peritumor. Sample size: Glioma Tumor (TCGA, n = 166), Glioma Peritumor (TCGA, n = 5), Normal Brain Cortex (GTEx, n = 105). **(C)** Validation of GTPase assay using varying concentrations of purified MIRO1 (GST-MIRO1). n=3. **(D)** MIRO1 protein was immunoprecipitated (IPed) from T98G cells using anti-MIRO1. The IPed MIRO1 protein was incubated with the indicated compound for 1 hour (10 μM), and GTPase activity was measured (amount of GTP hydrolyzed to free phosphate (μM) in each condition). n=4. All error bars represent the mean ± SD.

### Supplementary Tables

**Table S1. Glioma proteomics.**

**Table S2. Top hits from MIRO1 binder screens.**

**Table S3. *In vivo* pharmacokinetics of MB72 and MB08.**

### Supplementary Movie

**Movie S1. Cryo-ET tomogram and 3D segmentation of mitochondrial membrane remodeling upon BAX activation.** Sequential slices through a representative cryo-ET tomogram of a T98G cell treated with AAQ for 2 hours as in Fig. 6, showing localized distension of OMM from the inner boundary membrane. The 3D segmentation highlights the outer and inner mitochondrial membranes (blue), cristae (green), and adjacent vesicular organelles (red).

Fig. S1

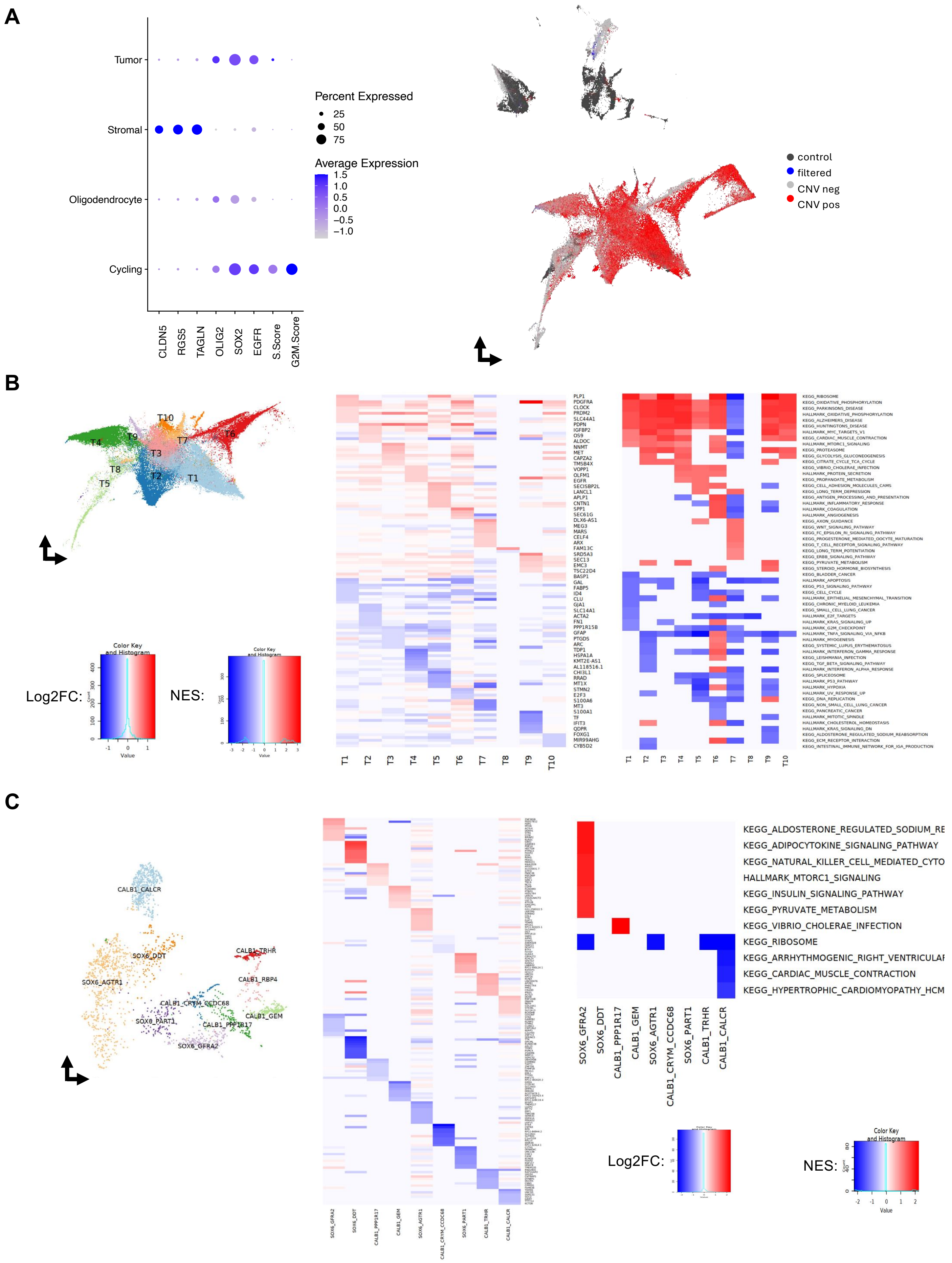
A
Tumor
Stromal
Oligodendrocyte
Cycling
CLDN5
RGS5
TAGLN
OLIG2
SOX2
EGFR
S.Score
G2M.Score
Percent Expressed
25
50
75
Average Expression
1.5
1.0
0.5
0.0
−0.5
−1.0
control
filtered
CNV neg
CNV pos
B
T10
T9
T4
T7
T6
T3
T8
T2
T1
T5
Color Key and Histogram
Log2FC:
NES:
Value
PLP1
PDGFRA
CLOCK
PRDM2
SLC44A1
PDPN
IGFBP2
OS9
ALDOC
NNMT
MET
CAPZA2
TMSB4X
VOPP1
OLFM1
EGFR
SEC14L2
LANCL1
APLP1
CNTN1
SPP1
SEC61G
DLX6-AS1
MEG3
MARS
CELF4
ARX
FAM13C
SRD5A3
SEC13
EMC3
TSC22D4
BASP1
GAL
FABP5
ID4
CLU
GJA1
SLC14A1
ACTA2
FN1
PPP1R15B
GFAP
PTGDS
ARC
TDP1
HSPA1A
KMT2E-AS1
AL118516.1
CHI3L1
RRAD
MT1X
STMN2
E2F3
S100A6
MT3
S100A1
TF
IFIT3
QDPR
FOXG1
MIR99AHG
CYB5D2
KEGG_RIBOSOME
KEGG_OXIDATIVE_PHOSPHORYLATION
KEGG_PARKINSONS_DISEASE
HALLMARK_OXIDATIVE_PHOSPHORYLATION
KEGG_ALZHEIMERS_DISEASE
KEGG_HUNTINGTONS_DISEASE
HALLMARK_MYC_TARGETS_V1
KEGG_CARDIAC_MUSCLE_CONTRACTION
HALLMARK_MTORC1_SIGNALING
KEGG_PROTEASOME
KEGG_GLYCOLYSIS_GLUCONEOGENESIS
KEGG_CITRATE_CYCLE_TCA_CYCLE
KEGG_VIBRIO_CHOLERAE_INFECTION
HALLMARK_PROTEIN_SECRETION
KEGG_PROPANOATE_METABOLISM
KEGG_CELL_ADHESION_MOLECULES_CAMS
KEGG_LONG_TERM_DEPRESSION
KEGG_ANTIGEN_PROCESSING_AND_PRESENTATION
HALLMARK_INFLAMMATORY_RESPONSE
HALLMARK_COAGULATION
HALLMARK_ANGIOGENESIS
KEGG_AXON_GUIDANCE
KEGG_WNT_SIGNALING_PATHWAY
KEGG_FC_EPSILON_RI_SIGNALING_PATHWAY
KEGG_PROGESTERONE_MEDIATED_OOCYTE_MATURATION
KEGG_T_CELL_RECEPTOR_SIGNALING_PATHWAY
KEGG_LONG_TERM_POTENTIATION
KEGG_ERBB_SIGNALING_PATHWAY
KEGG_PYRUVATE_METABOLISM
KEGG_STEROID_HORMONE_BIOSYNTHESIS
KEGG_BLADDER_CANCER
HALLMARK_APOPTOSIS
KEGG_P53_SIGNALING_PATHWAY
KEGG_CELL_CYCLE
HALLMARK_EPITHELIAL_MESENCHYMAL_TRANSITION
KEGG_CHRONIC_MYELOID_LEUKEMIA
KEGG_SMALL_CELL_LUNG_CANCER
HALLMARK_E2F_TARGETS
HALLMARK_KRAS_SIGNALING_UP
HALLMARK_G2M_CHECKPOINT
HALLMARK_TNFA_SIGNALING_VIA_NFKB
HALLMARK_MYOGENESIS
KEGG_SYSTEMIC_LUPUS_ERYTHEMATOSUS
HALLMARK_INTERFERON_GAMMA_RESPONSE
KEGG_LEISHMANIA_INFECTION
KEGG_TGF_BETA_SIGNALING_PATHWAY
HALLMARK_INTERFERON_ALPHA_RESPONSE
KEGG_SPLICEOSOME
HALLMARK_P53_PATHWAY
HALLMARK_HYPOXIA
HALLMARK_UV_RESPONSE_UP
KEGG_DNA_REPLICATION
KEGG_NON_SMALL_CELL_LUNG_CANCER
KEGG_PANCREATIC_CANCER
HALLMARK_MITOTIC_SPINDLE
HALLMARK_CHOLESTEROL_HOMEOSTASIS
HALLMARK_KRAS_SIGNALING_DN
KEGG_ALDOSTERONE_REGULATED_SODIUM_REABSORPTION
KEGG_ECM_RECEPTOR_INTERACTION
KEGG_INTESTINAL_IMMUNE_NETWORK_FOR_IGA_PRODUCTION
T1
T2
T3
T4
T5
T6
T7
T8
T9
T10
C
CALB1_CALCR
SOX6_DDT
CALB1_TRHR
SOX6_AGTR1
CALB1_RBP4
CALB1_CRYM_CCDC68
SOX6_PART1
CALB1_PPP1R17
CALB1_GEM
SOX6_GFRA2
KEGG_ALDOSTERONE_REGULATED_SODIUM_RE
KEGG_ADIPOCYTOKINE_SIGNALING_PATHWAY
KEGG_NATURAL_KILLER_CELL_MEDIATED_CYTO
HALLMARK_MTORC1_SIGNALING
KEGG_INSULIN_SIGNALING_PATHWAY
KEGG_PYRUVATE_METABOLISM
KEGG_VIBRIO_CHOLERAE_INFECTION
KEGG_RIBOSOME
KEGG_ARRHYTHMOGENIC_RIGHT_VENTRICULAI
KEGG_CARDIAC_MUSCLE_CONTRACTION
KEGG_HYPERTROPHIC_CARDIOMYOPATHY_HCM
SOX6_GFRA2
SOX6_DDT
CALB1_PPP1R17
CALB1_GEM
SOX6_AGTR1
CALB1_CRYM_CCDC68
SOX6_PART1
CALB1_TRHR
CALB1_CALCR
Log2FC:
NES:
Color Key and Histogram
Value

Fig. S2

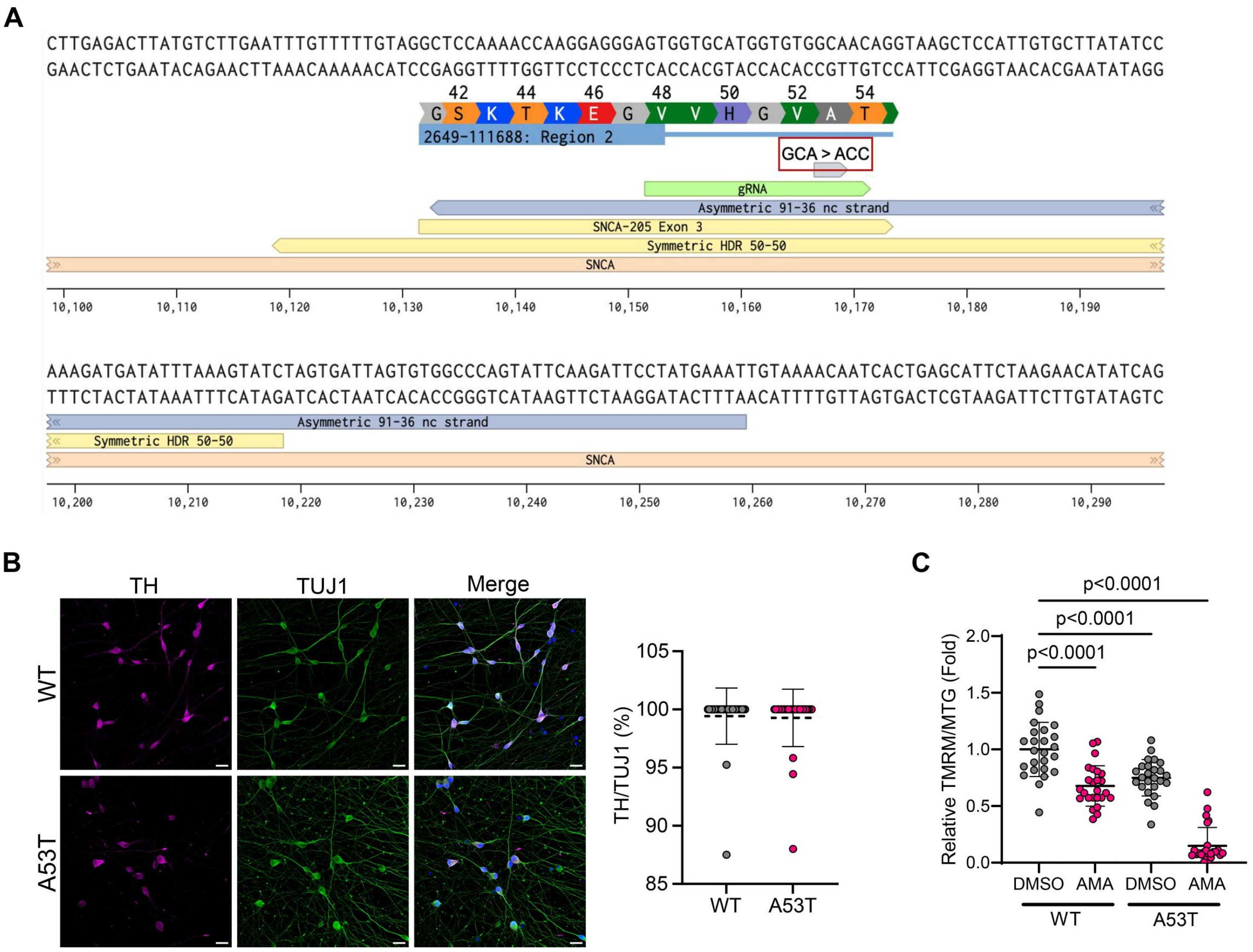
A
CTTGAGACTTATGTCTTGAATTTGTTTTTGTAGGCTCCAAAACCAAGGAGGGAGTGGTGCATGGTGTGGCAACAGGTAAGCTCCATTGTGCTTATATCC
GAACTCTGAATACAGAACTTAAACAAAAACATCCGAGGTTTTGGTTCCTCCCTCACCACGTACCACACCGTTGTCCATTCGAGGTAACACGAATATAGG
42 44 46 48 50 52 54
G S K T K E G V V H G V A T
2649-111688: Region 2
GCA > ACC
gRNA
Asymmetric 91-36 nc strand
SNCA-205 Exon 3
Symmetric HDR 50-50
SNCA
10,100 10,110 10,120 10,130 10,140 10,150 10,160 10,170 10,180 10,190
AAAGATGATATTTAAAGTATCTAGTGATTAGTGTGGCCCAGTATTCAAGATTCCTATGAAATTGTAAAACAATCACTGAGCATTCTAAGAACATATCAG
TTTCTACTATAAATTTCATAGATCACTAATCACACCGGGTCATAAGTTCTAAGGATACTTTAACATTTTGTTAGTGACTCGTAAGATTCTTGTATAGTC
Asymmetric 91-36 nc strand
Symmetric HDR 50-50
SNCA
10,200 10,210 10,220 10,230 10,240 10,250 10,260 10,270 10,280 10,290
B
TH
TUJ1
Merge
WT
A53T
TH/TUJ1 (%)
105
100
95
90
85
WT
A53T
C
p<0.0001
p<0.0001
p<0.0001
Relative TMRM/MTG (Fold)
2.0
1.5
1.0
0.5
0.0
DMSO
AMA
DMSO
AMA
WT
A53T

Fig. S3

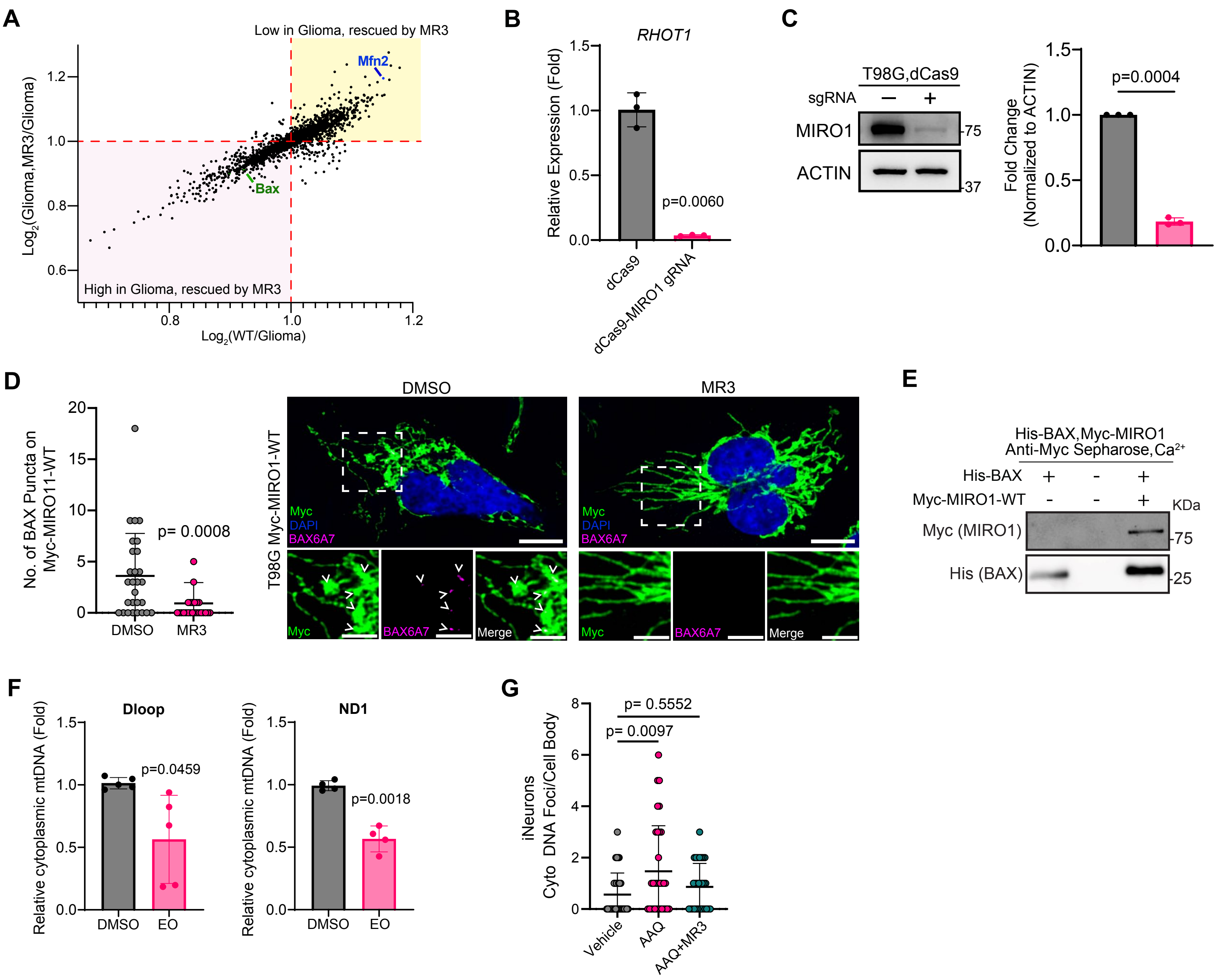

A
Low in Glioma, rescued by MR3
High in Glioma, rescued by MR3
Mfn2
Bax
Log2(Glioma,MR3/Glioma)
Log2(WT/Glioma)
B
RHOT1
Relative Expression (Fold)
p=0.0060
dCas9
dCas9-MIRO1 gRNA
C
T98G,dCas9
sgRNA
MIRO1
ACTIN
75
37
Fold Change (Normalized to ACTIN)
p=0.0004
D
No. of BAX Puncta on Myc-MIRO11-WT
p= 0.0008
DMSO
MR3
T98G Myc-MIRO1-WT
Myc
DAPI
BAX6A7
Merge
E
His-BAX,Myc-MIRO1
Anti-Myc Sepharose,Ca2+
His-BAX
Myc-MIRO1-WT
KDa
Myc (MIRO1)
His (BAX)
75
25
F
Dloop
ND1
Relative cytoplasmic mtDNA (Fold)
p=0.0459
p=0.0018
DMSO
EO
G
iNeurons Cyto DNA Foci/Cell Body
p= 0.5552
p= 0.0097
Vehicle
AAQ
AAQ+MR3

Fig. S4

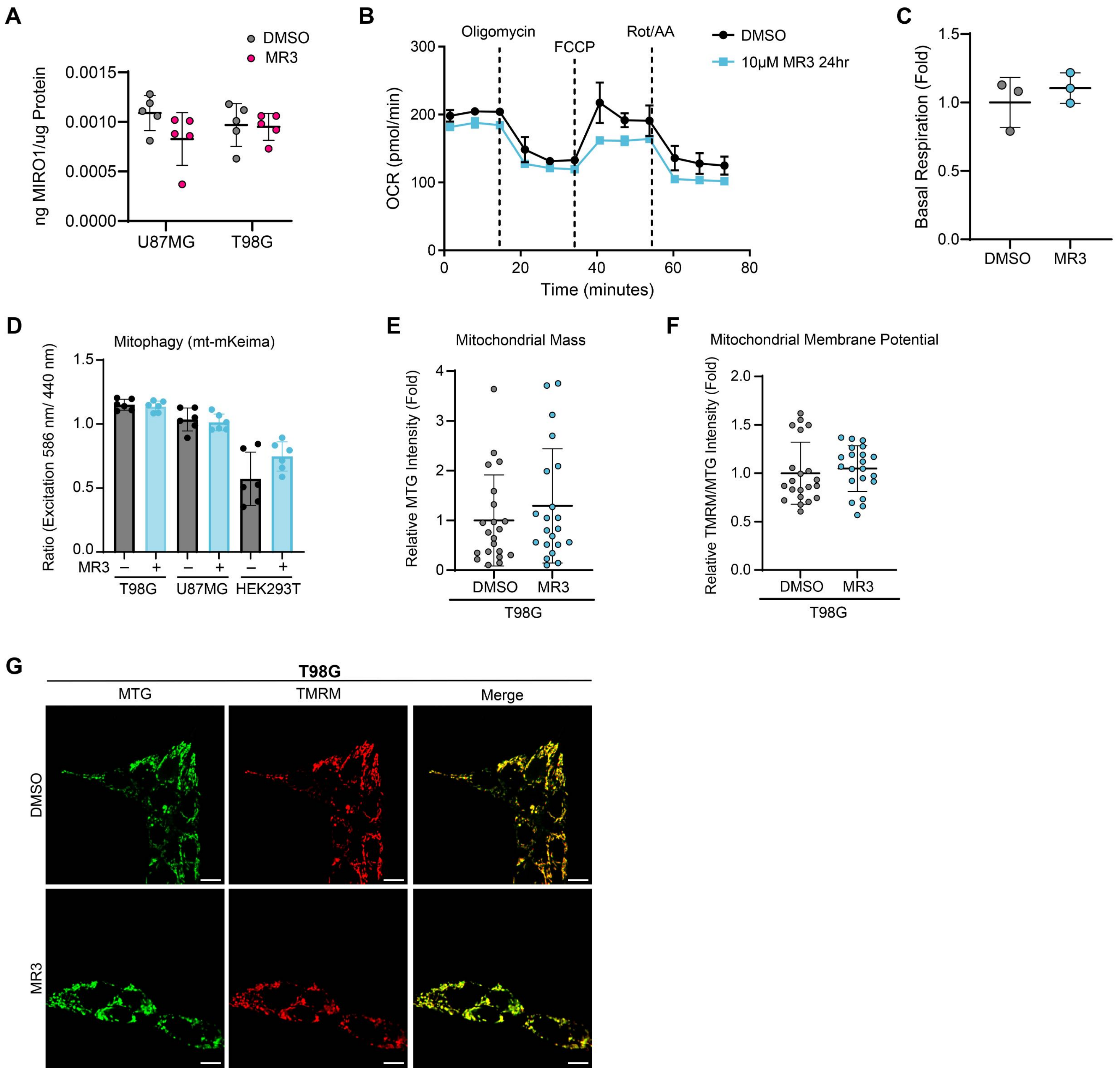
A
DMSO
MR3
ng MIRO1/ug Protein
0.0015
0.0010
0.0005
0.0000
U87MG
T98G
B
Oligomycin
FCCP
Rot/AA
DMSO
10μM MR3 24hr
OCR (pmol/min)
300
200
100
0
0
20
40
60
80
Time (minutes)
C
Basal Respiration (Fold)
1.5
1.0
0.5
0.0
DMSO
MR3
D
Mitophagy (mt-mKeima)
Ratio (Excitation 586 nm/ 440 nm)
1.5
1.0
0.5
0.0
MR3
–
+
T98G
U87MG
HEK293T
E
Mitochondrial Mass
Relative MTG Intensity (Fold)
4
3
2
1
0
DMSO
MR3
T98G
F
Mitochondrial Membrane Potential
Relative TMRM/MTG Intensity (Fold)
2.0
1.5
1.0
0.5
0.0
DMSO
MR3
T98G
G
T98G
MTG
TMRM
Merge
DMSO
MR3

Fig. S5

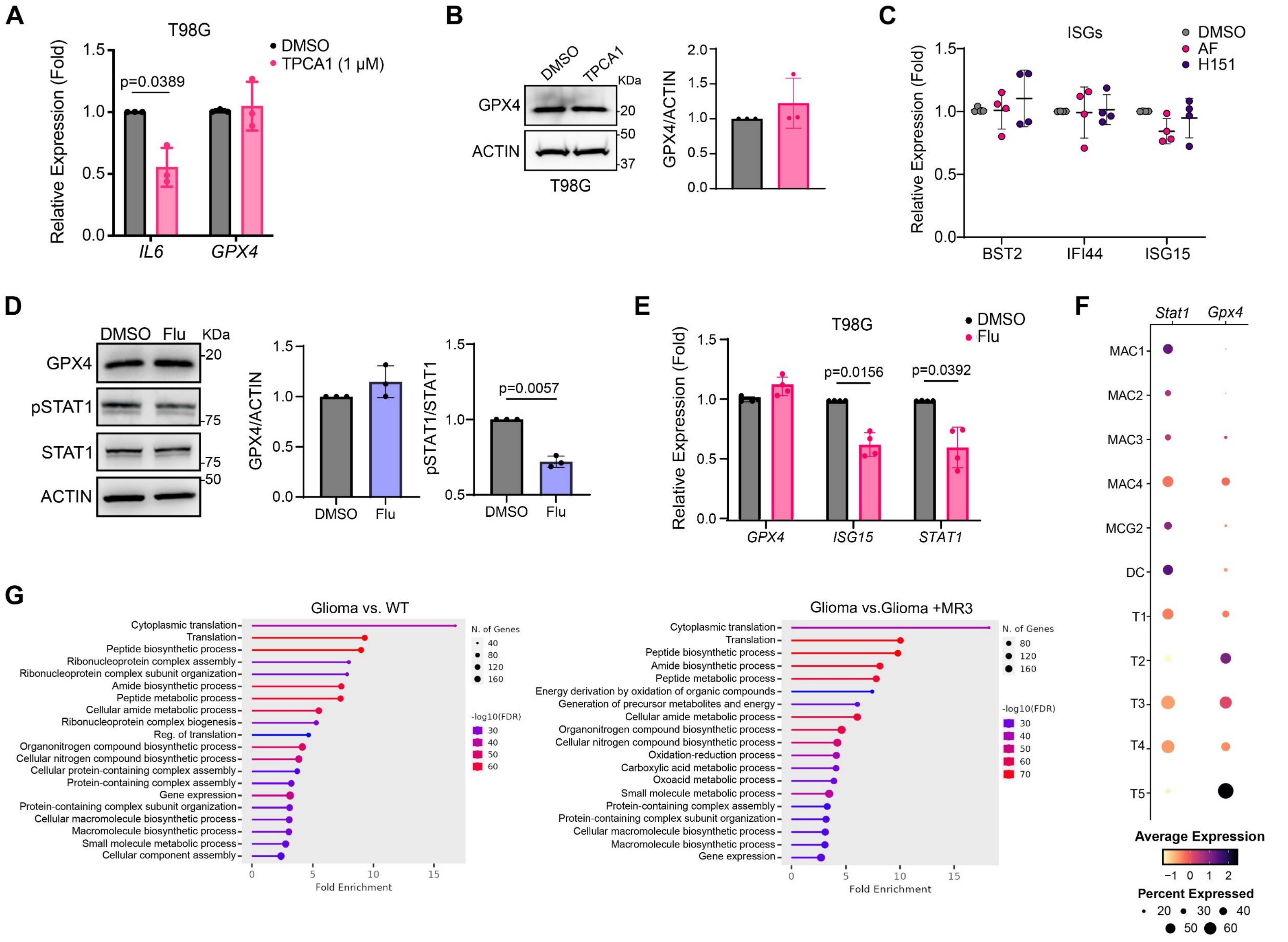

A
T98G
DMSO
TPCA1 (1 μM)
p=0.0389
Relative Expression (Fold)
IL6
GPX4
B
DMSO
TPCA1
KDa
GPX4
ACTIN
T98G
GPX4/ACTIN
C
ISGs
DMSO
AF
H151
Relative Expression (Fold)
BST2
IFI44
ISG15
D
DMSO
Flu
KDa
GPX4
pSTAT1
STAT1
ACTIN
GPX4/ACTIN
pSTAT1/STAT1
p=0.0057
E
T98G
DMSO
Flu
p=0.0156
p=0.0392
Relative Expression (Fold)
GPX4
ISG15
STAT1
F
Stat1
Gpx4
MAC1
MAC2
MAC3
MAC4
MCG2
DC
T1
T2
T3
T4
T5
Average Expression
Percent Expressed
G
Glioma vs. WT
Cytoplasmic translation
Translation
Peptide biosynthetic process
Ribonucleoprotein complex assembly
Ribonucleoprotein complex subunit organization
Amide biosynthetic process
Peptide metabolic process
Cellular amide metabolic process
Ribonucleoprotein complex biogenesis
Reg. of translation
Organonitrogen compound biosynthetic process
Cellular nitrogen compound biosynthetic process
Cellular protein-containing complex assembly
Protein-containing complex assembly
Gene expression
Protein-containing complex subunit organization
Cellular macromolecule biosynthetic process
Macromolecule biosynthetic process
Small molecule metabolic process
Cellular component assembly
Fold Enrichment
N. of Genes
-log10(FDR)
Glioma vs.Glioma +MR3
Cytoplasmic translation
Translation
Peptide biosynthetic process
Amide biosynthetic process
Peptide metabolic process
Energy derivation by oxidation of organic compounds
Generation of precursor metabolites and energy
Cellular amide metabolic process
Organonitrogen compound biosynthetic process
Cellular nitrogen compound biosynthetic process
Oxidation-reduction process
Carboxylic acid metabolic process
Oxoacid metabolic process
Small molecule metabolic process
Protein-containing complex assembly
Protein-containing complex subunit organization
Cellular macromolecule biosynthetic process
Macromolecule biosynthetic process
Gene expression
Fold Enrichment
N. of Genes
-log10(FDR)

Fig. S6

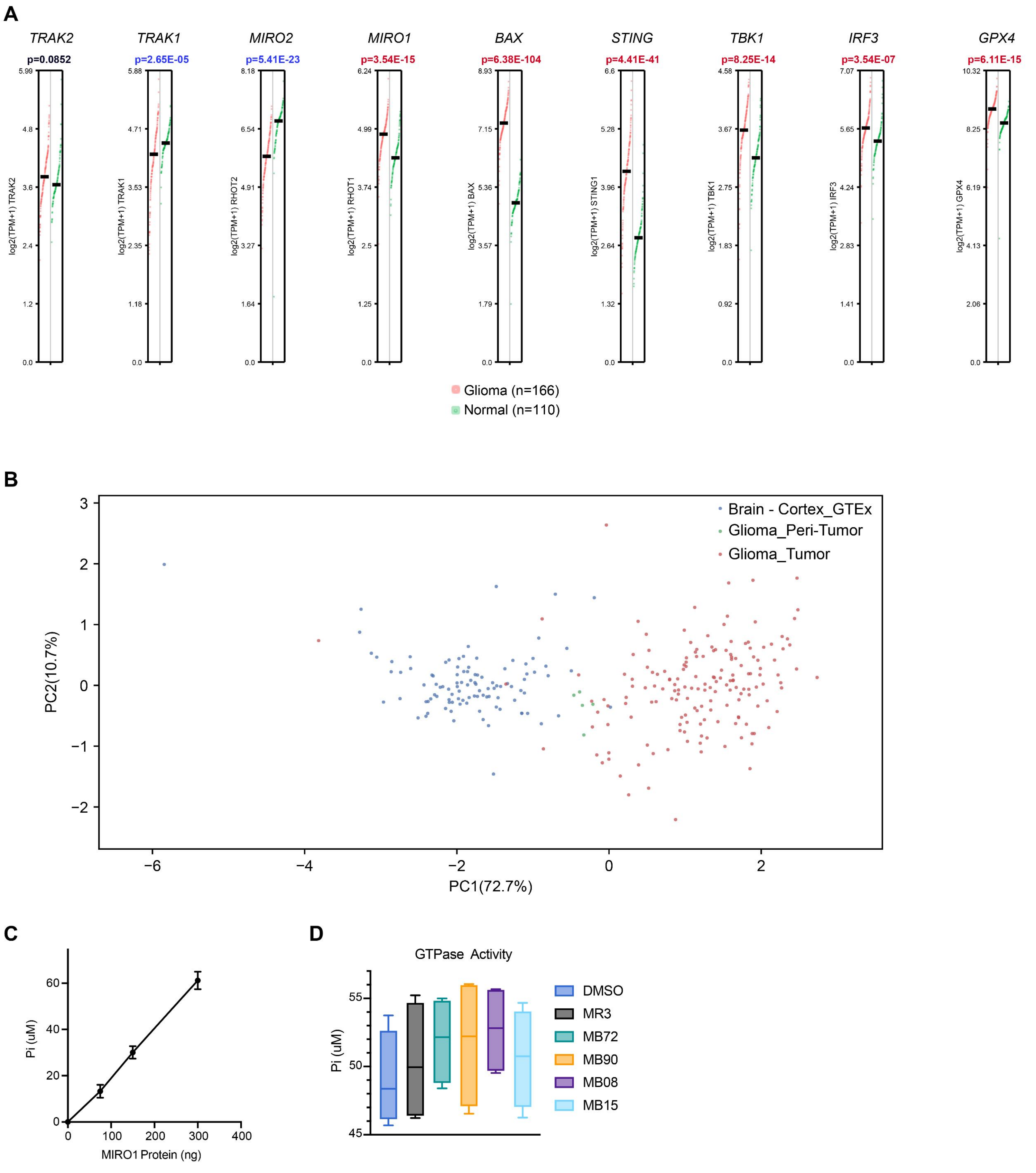

A
TRAK2
p=0.0852
log2(TPM+1) TRAK2
TRAK1
p=2.65E-05
log2(TPM+1) TRAK1
MIRO2
p=5.41E-23
log2(TPM+1) RHOT2
MIRO1
p=3.54E-15
log2(TPM+1) RHOT1
BAX
p=6.38E-104
log2(TPM+1) BAX
STING
p=4.41E-41
log2(TPM+1) STING1
TBK1
p=8.25E-14
log2(TPM+1) TBK1
IRF3
p=3.54E-07
log2(TPM+1) IRF3
GPX4
p=6.11E-15
log2(TPM+1) GPX4
Glioma (n=166)
Normal (n=110)
B
Brain - Cortex_GTEx
Glioma_Peri-Tumor
Glioma_Tumor
PC2(10.7%)
PC1(72.7%)
C
Pi (uM)
MIRO1 Protein (ng)
D
GTPase Activity
Pi (uM)
DMSO
MR3
MB72
MB90
MB08
MB15
